# Engineering *S. cerevisiae* extracellular vesicles using synthetic biology

**DOI:** 10.64898/2026.03.06.710173

**Authors:** Jeff Bouffard, Joseph Trani, Alexander Christophe Pawelczak, Marie Laufens, Mόnica Núñez-Soto, Christopher Leonard Brett

**Affiliations:** Department of Biology, Concordia University, 7141 Sherbrooke St. W., Montreal, Quebec, Canada, H4B 1R6

**Keywords:** Extracellular vesicle, exosome, *Saccharomyces cerevisiae*, synthetic biology, genetic engineering, drug delivery, Bro1, CD63, PDGFR

## Abstract

Extracellular vesicles (EVs) hold great promise as therapeutic delivery vehicles, leveraging their natural role as mediators of intercellular communication in all organisms studied. However, many barriers must be overcome to realize their full potential. *Saccharomyces cerevisiae* is an attractive chassis organism to explore solutions: It is used for drug biomanufacturing, it is amenable to complex genetic engineering, and their EVs can drive responses in human cells. To further develop this prospect, we sought to genetically modify *S. cerevisiae* EVs by devising a research framework amenable to iterative design, build, test, learn cycles – a core principle of synthetic biology. Using this approach, we focused on identifying new scaffolds – proteins that load cargoes into EVs – from a small pool of candidates. We first optimized a modular cloning strategy, called “EVclo”, for plasmid and genome-integrated candidate gene expression.

Candidate genes were fused to EGFP, and after confirming expression in cells, we showed that scaffold-EFGP proteins colocalized with mRuby2-tagged Nhx1, a biomarker of multivesicular bodies, presumed sites of EV biogenesis. We triggered release of EVs by heat stress, isolated these EVs by ultrafiltration and size exclusion chromatography, and confirmed the presence of exosome-sized EVs in all samples. We find that candidate scaffold proteins did not affect EV size, morphology or titers. Further analysis of these samples indicated that some EGFP-tagged scaffolds are present in EVs: Bro1, a yeast ortholog of ALIX, was most abundant and ExoSignal showed highest enrichment of the human candidates. In all, we conclude that Bro1 is a good scaffold for future engineering strategies, and that human proteins can be sorted into yeast EVs suggesting conservation of the sorting machinery and demonstrating that yeast EVs can be humanized. This synthetic biology-based, proof-of-concept study establishes *S. cerevisiae* as a platform to engineer and bioproduce designer EVs for many applications.

**GRAPHICAL ABSTRACT:** 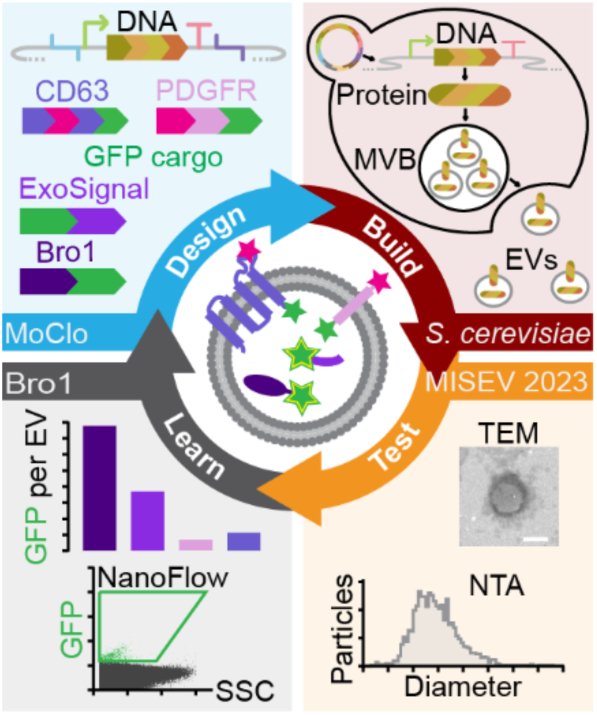

**HIGHLIGHTS AND TOC BLURB:**

- synthetic biology-based system was optimized to engineer EVs in *S. cerevisiae*
- EV scaffolds can be sorted to yeast EVs
- is an efficient scaffold to sort proteins into yeast EVs
- *S. cerevisiae* can be used to engineer designer EVs for drug delivery

Extracellular vesicles (EVs) are a promising new modality for drug delivery. However, designer EVs must be engineered to broaden applications and improve efficacy. Here, Bouffard et al. optimize methods rooted in synthetic biology to genetically engineer EVs in *S. cerevisiae*, a yeast commonly used to manufacture biological drugs. They find that ectopically expressed human EV scaffolds (CD63, ExoSignal, PDGFR) can be sorted to yeast EVs, but Bro1 – the yeast ortholog of ALIX – was most efficient at sorting GFP into EVs. This proof-of-concept study demonstrates a single DBTL (design-build-test-learn) cycle that can be used to develop designer EVs for therapeutic applications.

## 1 | INTRODUCTION

Thought to mediate intercellular communication and/or waste clearance, extracellular vesicles (EVs) are natural lipid nanoparticles (LNPs) released by every organism studied – from bacteria to yeast to humans (Gill et al., 2019) (Abels & Breakefield, 2016) (Ludwig & Giebel, 2012) (van Niel et al., 2022) (Yáñez-Mó et al., 2015) (Tkach & Théry, 2016) (Welsh et al., 2024). For communication, donor cells are thought to selectively load EVs with bioactive cargoes (proteins, lipids, nucleic acids) either displayed on the outer surface of their lipid bilayer, imbedded in this membrane, or contained within the lumen (Colombo et al., 2014) (Kalluri & LeBleu, 2020).

After release into the surrounding environment (e.g. interstitial fluid, blood, urine, saliva, cerebral spinal fluid), EVs are selectively recognized by target cells and deliver their cargoes (Mathieu et al., 2019) to elicit physiological responses underlying wound healing, reproduction and tissue development, among others in mammals (Yáñez-Mó et al., 2015). They are also implicated in the pathogenesis of cancers and neurodegenerative disorders (Amin et al., 2024) as well as in host–pathogen interactions underlying infection by playing roles in virulence and immune defense (Rodrigues et al., 2018). As such, pathogenic EV components show great promise as body fluid analytes for improved diagnostics. And as they represent nature’s LNPs with many desirable innate properties, EVs show immense potential to become a novel drug delivery modality that could transform medicine (Kumar et al., 2024).

However, to maximally realize possible therapeutic applications, EVs must be modified through engineering (Jafari et al., 2020). This involves strategies for loading desired proteins, RNAs, DNAs or small molecules into the lumen of EVs to precisely modulate behaviors of target cells (Danilushkina et al., 2023) (Erana-Perez et al., 2024). Also, customized moieties on EV surfaces can be added to enhance stability (e.g. increase retention time in the body by preventing clearance or degradation), to accurately target specific tissues or cell types, to trigger receptor-mediated signaling that drives therapeutic responses, and/or to enhance uptake by target cells.

Both strategies may involve exogenous methods by directly modifying purified EVs, or endogenous methods during EV biogenesis within cells through genetic engineering. Although powerful, the latter is largely underdeveloped due to many challenges, including an incomplete understanding of the mechanisms responsible for EV biogenesis. However, one approach is to fuse therapeutic protein cargoes to “scaffold” proteins that are intrinsically sorted into EVs.

Examples include the mammalian EV biomarker CD63, a tetraspanin polytopic protein embedded in small EV membranes that can accept cargo fusions to its surface or lumenal faces, and the small polypeptide ExoSignal that can direct cargoes to the EV lumen (Stickney et al., 2016) (Ferreira et al., 2022).

Primarily derived from human mesenchymal stem cells, engineered EVs for treatment of cancers, diabetes, cardiovascular disease and neurological disorders (Mentkowski et al., 2018) show promise with some clinical trials in progress (Herrmann et al., 2021) (Van Delen et al., 2024) (Mizenko et al., 2024) (Ma et al., 2025). Despite these advancements, many barriers must be overcome to fully realize EV-mediated drug delivery in the clinic, e.g. producing sufficient EVs for widespread use, optimizing methods to purify desired EVs from heterogenous samples, and improving cargo loading, cell targeting and cellular uptake for better efficacy (Paganini et al., 2019) (Adlerz et al., 2020) (Estes et al., 2022) (Ng et al., 2022) (Xu et al., 2025). Given that EVs are released from all organisms studied, spanning the entire tree of life (Gill et al., 2019) (Woith et al., 2019) (Liebana-Jordan et al., 2021), scientists are considering alternative, non-human platforms to develop therapeutic EVs while addressing current limitations (Oliveira et al., 2010) (Zhao et al., 2019). Because it is currently used for large-scale biomanufacturing of drugs (Buchholz & Collins, 2013), the Generally Regarded As Safe (GRAS) yeast *Saccharomyces cerevisiae* (or “baker’s yeast”) and its EVs are being considered as vaccine adjuvants (Higuchi et al., 2023), delivery vehicles for therapeutic RNAs (Yuan et al., 2024), and advanced probiotics (Gryciuk et al., 2025). In support, work from our lab demonstrates that *S. cerevisiae* releases a relatively homogenous population of EVs in response to mild heat stress (Logan et al., 2024), and *S. cerevisiae* is notoriously amenable to complex genetic engineering, making it an ideal platform to develop and produce engineered EVs for therapeutic applications.

Specifically, *S. cerevisiae* is a common chassis organism used to support the powerful and emerging field of synthetic biology (Endy, 2005) (Garner, 2021), which includes established research frameworks and tools that can be leveraged to expedite engineering of designer EVs (Kelwick et al., 2024). One core principle of synthetic biology is abstraction to modularity (Garcia & Trinh, 2019) that entails viewing genetic sequences as modular parts – e.g. promoters, coding sequences (CDS), terminators – fused in combination using a common DNA assembly standard to build complex genetic circuits (Lee et al., 2015) (Malcl et al., 2022). Another core principle is the use of the Design-Build-Test-Learn (DBTL) cycle (Whitford et al., 2021) (Jeon et al., 2025), shorthand for an engineering design process that provides a systematic, iterative framework for efficient and successful project advancement (Lawson et al., 2019). Many EV engineering efforts implicitly consider genetic modularity and perform DBTL-like cycles to iteratively improve their designer EVs (Kojima et al., 2018) (Dooley et al., 2021) (Stranford et al., 2024), but without explicitly using a synthetic biology framework that will increase productivity (Kelwick et al., 2024) (David et al., 2021) (Freemont, 2019).

Here, we report a proof-of-concept study focused on engineering designer EVs in *S. cerevisiae* using a synthetic biology framework, with hopes of realizing their potential for human therapeutics and other applications. We designed, built and tested genetic constructs to engineer EVs using popular human scaffolds and a promising yeast scaffold, domesticating the parts to conform to the widely used Yeast Tool Kit (YTK) MoClo assembly standard (Lee et al., 2015) and to accommodate markerless genomic integration using the MyLO toolkit (Bean et al., 2022). After a single DBTL cycle, we established a workflow and identified useful EV scaffolds that will support further development of *S. cerevisiae* as a platform for the engineering and bioproduction of therapeutic EVs.

## 2 | METHODS

### 2.1 | Molecular cloning

All genetic sequences were designed, manipulated, and analyzed using Geneious Prime software, version 2024.0.5, or previous versions. Constructs in Figure 1 are represented using the Synthetic Biology Open Language (SBOL) Visual standard version 3.0 (Baig et al., 2021), although the SBOL data model was not used. Table 1 provides primers, Table 2 provides plasmids, and Table 3 provides the sequences of EVclo scaffold CDS parts used in this study.

**Figure 1.**
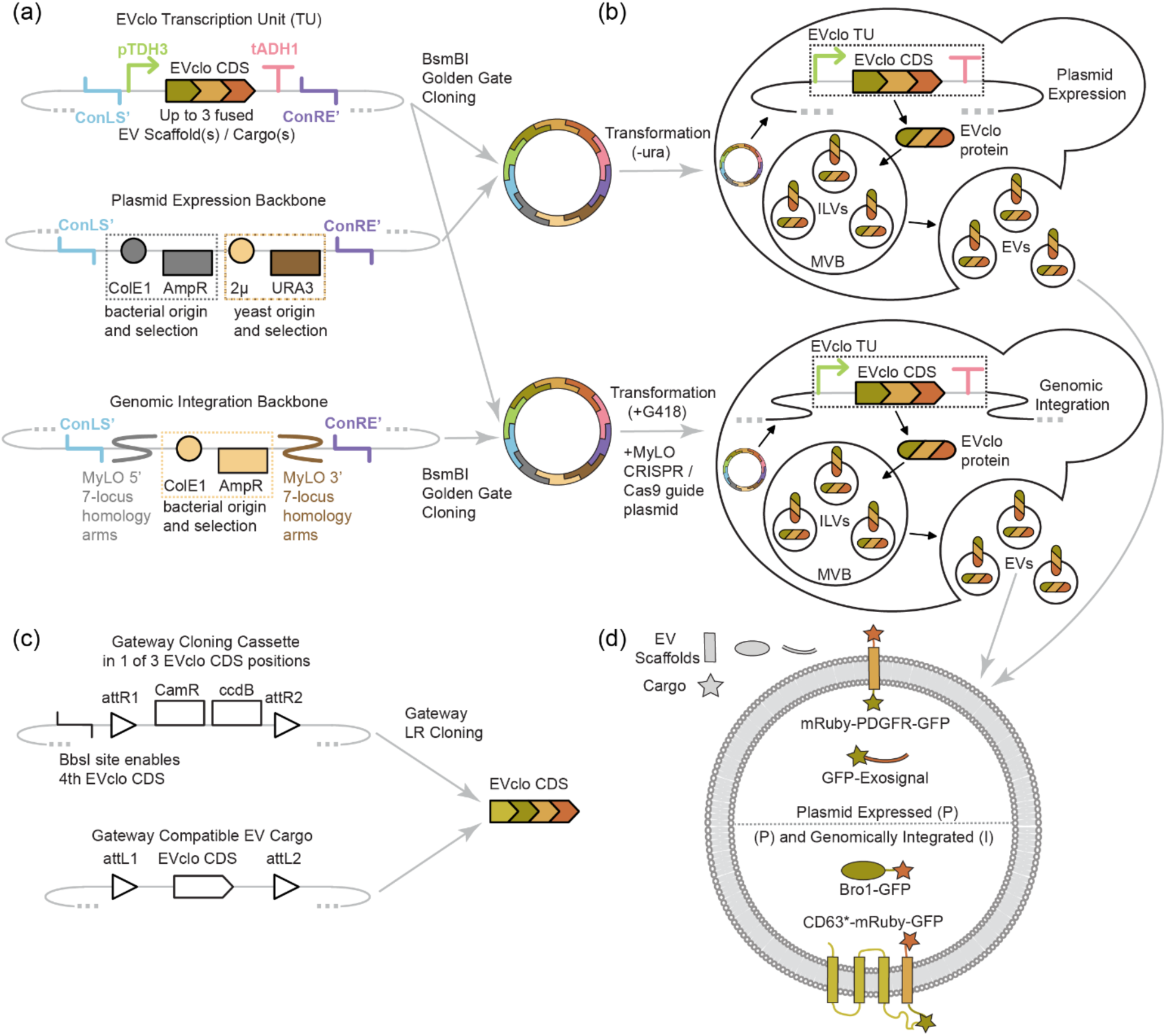
EVclo: A synthetic biology approach to engineering EVs in *S. cerevisiae*. (a) Schematic of modular genetic constructs. Parts follow the Yeast Toolkit (YTK) assembly standard to generate plasmids via BsaI Golden Gate reactions. EVclo transcription units (TU) (top) are composed of a strong constitutive promoter (pTDH3), EVclo coding sequences (CDS) featuring 2 or 3 EV scaffolds and cargos as a single translational fusion, and a terminator (tADH1). Connector parts (ConLS’, ConRE’) allow TU relocation between vectors using BsmBI Golden Gate reactions. For plasmid-based expression (middle) of EVclo TUs, the high copy number yeast origin of replication (2µ) was used, with the URA3 module providing auxotrophic selection. For genomic integration (bottom) of EVclo TUs, parts from the Markerless Yeast Localization and Overexpression (MyLO) toolkit featuring homology arms for 7 different loci in the yeast genome were used. Both backbones feature ConLS’ and ConRE’ connector parts, as well as the ColE1 bacterial origin of replication and AmpR ampicillin resistance module to propagate plasmids in *E.coli*. (b) Schematic of designer EV production in *S. cerevisiae*. EVclo TUs are introduced into yeast cells either on plasmids (top) or integrated into the genome (bottom). Yeast cells with EVclo TUs express EVclo proteins composed of EV scaffolds and cargos. Successful EV scaffolds associate with intralumenal vesicles (ILVs) within the multivesicular body (MVB), the site of EV biogenesis. When the MVB fuses with the plasma membrane, ILVs containing EVclo proteins are released, becoming designer EVs. (c) To enable screening libraries of EV scaffolds or cargos, Gateway cloning cassettes (top) were built as CDS type parts for inclusion in EVclo CDSs. Gateway cloning cassettes are composed of the attR1 and attR2 recombination sites, flanking a CamR chloramphenicol resistance module for selection and a ccdB bacterial toxin module to increase efficiency of Gateway cloning reactions. When combined with Gateway Entry clones (bottom) featuring an EVclo CDS flanked by attL1 and attL2 recombination sites in a Gateway LR reaction, the EVclo CDS replaces the CamR and ccdB modules, in frame with flanking EVclo CDSs at other positions. To enable the addition of a fourth EVclo CDS component, some Gateway cloning cassettes were built with a 5’ BbsI site. (d) Schematic of EVclo designer EVs tested in this study. Each engineered yeast strain produced only 1 of the scaffold proteins shown. The constructs in the top half were only tested using plasmid expression, the constructs in the bottom half were tested using both plasmid expression and genomic integration.

**Table 1 –.**
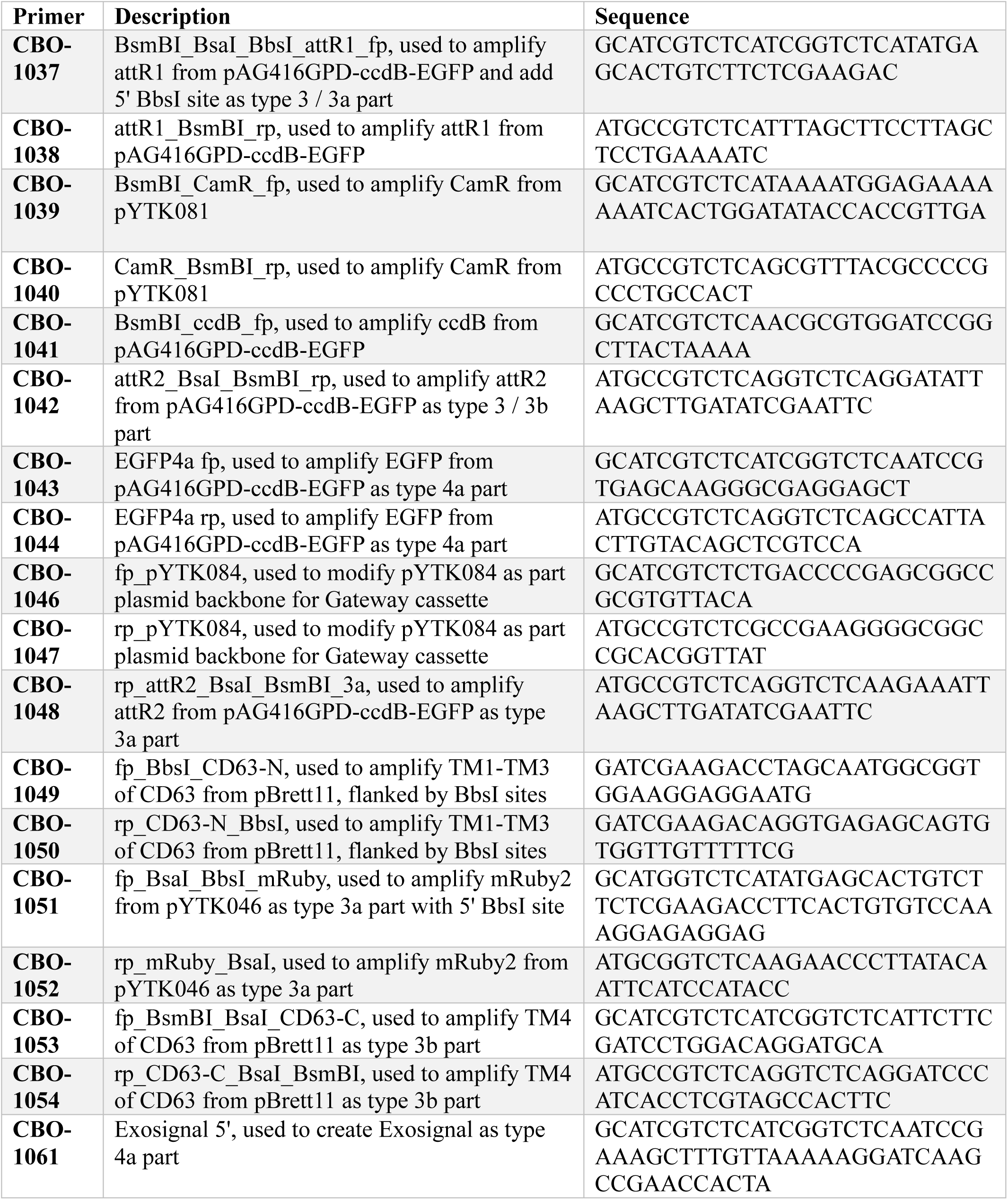
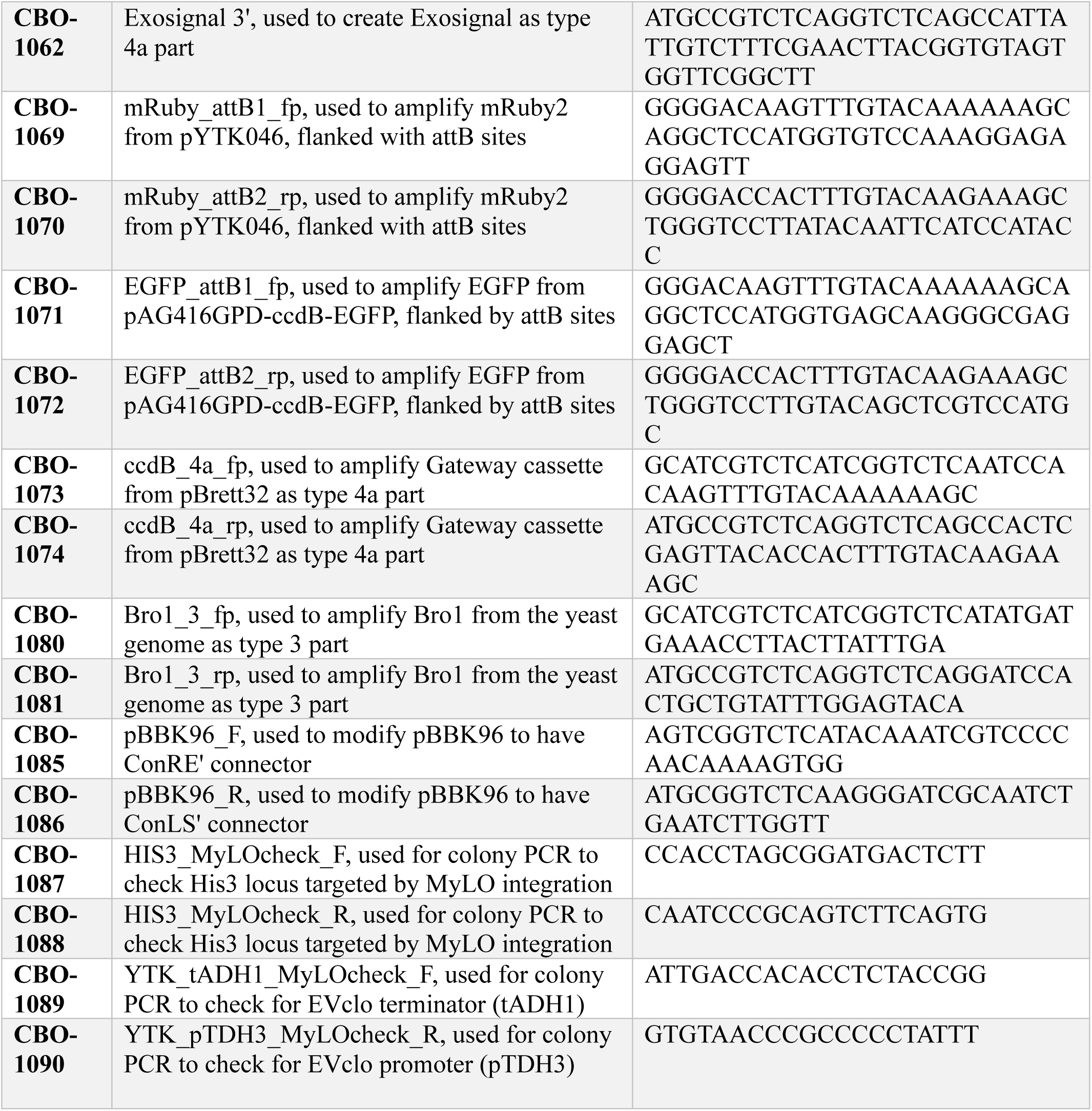
Primers used in this study.

**Table 2 –.** Plasmids used in this study.

| Plasmid | Description | Source |
| --- | --- | --- |
| **pAG416GPD-ccdB-EGFP** | source of attR sites and ccdB gene cassette for Gateway cassette parts, and EGFP parts and Entry clone | Addgene plasmid #14196 (Alberti et al., 2007) |
| **pBBK11** | MyLO Cas9 SplitKanR guide plasmid targeting FF18 locus | Addgene plasmid #178995 (Bean et al., 2022) |
| **pBBK17** | MyLO Cas9 SplitKanR guide plasmid targeting HIS3 locus | Addgene plasmid #179001 (Bean et al., 2022) |
| **pBBK96** | MyLO integration vector | Addgene plasmid #179080 (Bean et al., 2022) |
| **pBrett11** | Full length *H.sapiens* codon CD63 Expression clone | This study |
| **pBrett12** | Nhx1 Entry clone in pDONR221 backbone | This study |
| **pBrett25** | EVclo plasmid expression backbone assembled from pYTK008, 047, 073, 074, 082, and 083 | This study |
| **pBrett27** | GatewayCassette-mRuby2 Destination vector assembled from pBrett25, pYTK010, pBrett35, pYTK046, and pYTK053 | This study |
| **pBrett32** | Gateway cassette with 5' BbsI site type 3 part plasmid in pYTK084 backbone | This study |
| **pBrett35** | Gateway cassette with 5' BbsI site type 3a part plasmid in pYTK084 backbone | This study |
| **pBrett36** | EGFP type 4a part plasmid in pYTK001 backbone | This study |
| **pBrett38** | mRuby2-CD63(TM4)-EGFP Expression clone assembled from pBrett25, pYTK009, mRuby2 with 5' BbsI site type 3a fragment, CD63(TM4) type 3b fragment, pBrett36, and pYTK063 | This study |
| **pBrett40 \*** | CD63*-mRuby-EGFP Expression clone assembled from pBrett38 and CD63(TM1-3) BbsI fragment | This study |
| **pBrett65** | GatewayCassette-Exosignal Destination vector assembled from pBrett25, pYTK009, pBrett32, pBrett69, and pYTK063 | This study |
| **pBrett66** | PDGFR type 3b part plasmid in pYTK001 backbone | This study |
| **pBrett69** | Exosignal type 4a part plasmid in pYTK001 backbone | This study |
| **pBrett70 \*** | GFP-Exosignal Expression clone assembled by Gateway LR cloning with pBrett65 and pBrett91 | This study |
| **pBrett75** | Nhx1-mRuby Expression clone assembled by Gateway LR cloning of pBrett12 and pBrett27 | This study |
| **pBrett88** | Gateway cassette type 4a part plasmid in pYTK084 backbone | This study |
| **pBrett89** | GatewayCassette-PDGFR-EGFP Destination vector assembled from pBrett25, pYTK009, pBrett35, pBrett66, pBrett36, and pYTK063 | This study |
| **pBrett90** | mRuby2 Entry clone in pDONR221 backbone | This study |
| **pBrett91** | EGFP Entry clone in pDONR221 backbone | This study |
| **pBrett93 \*** | mRuby-PDGFR-EGFP Expression clone assembled by Gateway LR cloning of pBrett89 and pBrett90 | This study |
| **pBrett94** | Bro1 type 3 part plasmid in pYTK001 backbone | This study |
| **pBrett96** | Bro1-GatewayCassette Destination vector assembled using pBrett25, pYTK009, pBrett94, pBrett88, and pYTK063 | This study |
| **pBrett104** | pBBK96 modified to have ConLS' and ConRE' assembly connectors | This study |
| **pBrett107 \*** | Bro1-GFP Integration vector assembled by BsmBI reaction of pBrett104 and pBrett172 | This study |
| **pBrett108 \*** | CD63*-mRuby-GFP Integration vector assembled by BsmBI reaction of pBrett40 and pBrett104 | This study |
| **pBrett127 \*** | Nhx1-mRuby Integration vector assembled by BsmBI reaction of pBrett75 and pBrett104 | This study |
| **pBrett172 \*** | Bro1-GFP Expression clone assembled by Gateway LR cloning of pBrett91 and pBrett96 | This study |
| **pDONR221** | Empty Gateway Donor vector for making Entry clones | Invitrogen Corp. |
| **pYTK001** | Empty part plasmid | Addgene plasmid #65108 (Lee et al., 2015) |
| **pYTK008** | ConLS' part plasmid | Addgene plasmid #65115 (Lee et al., 2015) |
| **pYTK009** | pTDH3 promoter part plasmid | Addgene plasmid #65116 (Lee et al., 2015) |
| **pYTK010** | pCCW12 promoter part plasmid | Addgene plasmid #65117 (Lee et al., 2015) |
| **pYTK046** | source of mRuby2 parts and Entry clone in pDONR221 | Addgene plasmid #65153 (Lee et al., 2015) |
| **pYTK047** | superfolder GFP dropout cassette part plasmid | Addgene plasmid #65154 (Lee et al., 2015) |
| **pYTK053** | tADH1 terminator type 4 part plasmid | Addgene plasmid #65160 (Lee et al., 2015) |
| **pYTK063** | tADH1 terminator type 4b part plasmid | Addgene plasmid #65170 (Lee et al., 2015) |
| **pYTK073** | ConRE' part plasmid | Addgene plasmid #65180 (Lee et al., 2015) |
| **pYTK074** | URA3 yeast selection marker part plasmid | Addgene plasmid #65181 (Lee et al., 2015) |
| **pYTK081** | source of CamR gene for Gateway cassette parts | Addgene plasmid #65188 (Lee et al., 2015) |
| **pYTK082** | 2μ yeast origin of replication part plasmid | Addgene plasmid #65189 (Lee et al., 2015) |
| **pYTK083** | *E. coli* AmpR selection marker and Cole1 origin of replication part plasmid | Addgene plasmid #65190 (Lee et al., 2015) |
| **pYTK084** | modified to serve as part plasmid backbone for Gateway cassette parts | Addgene plasmid #65191 (Lee et al., 2015) |

**Table 3 –.**
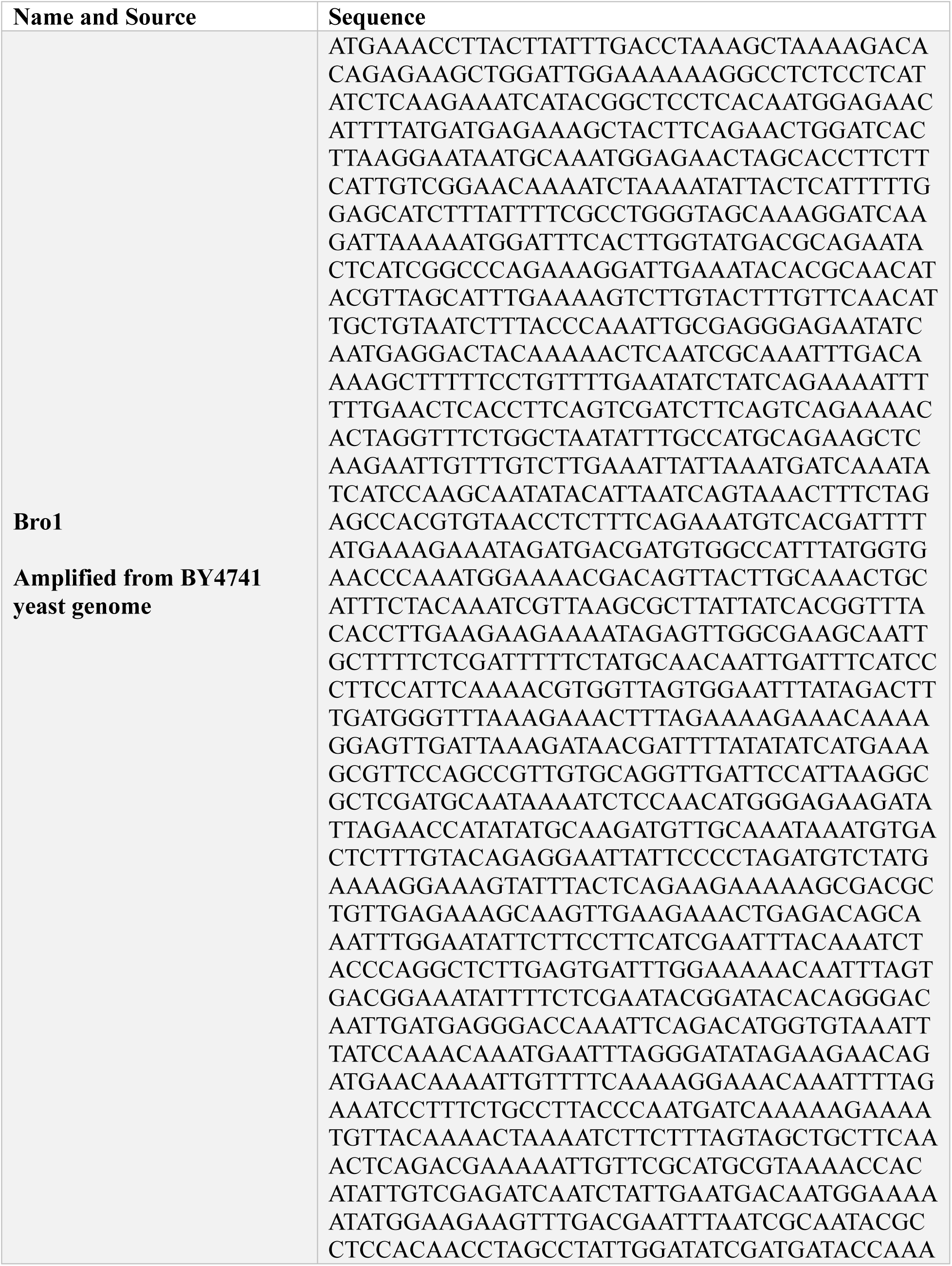
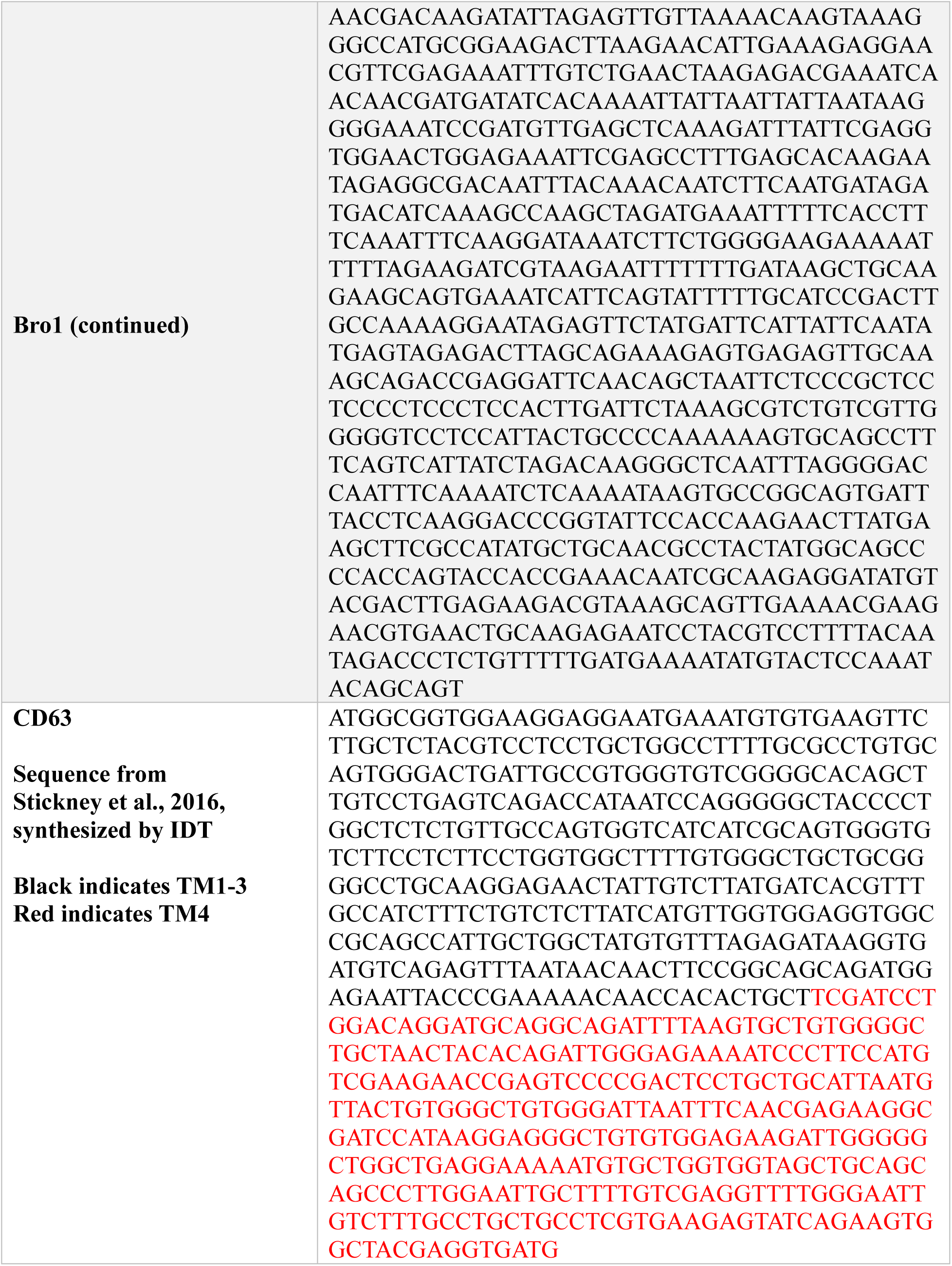
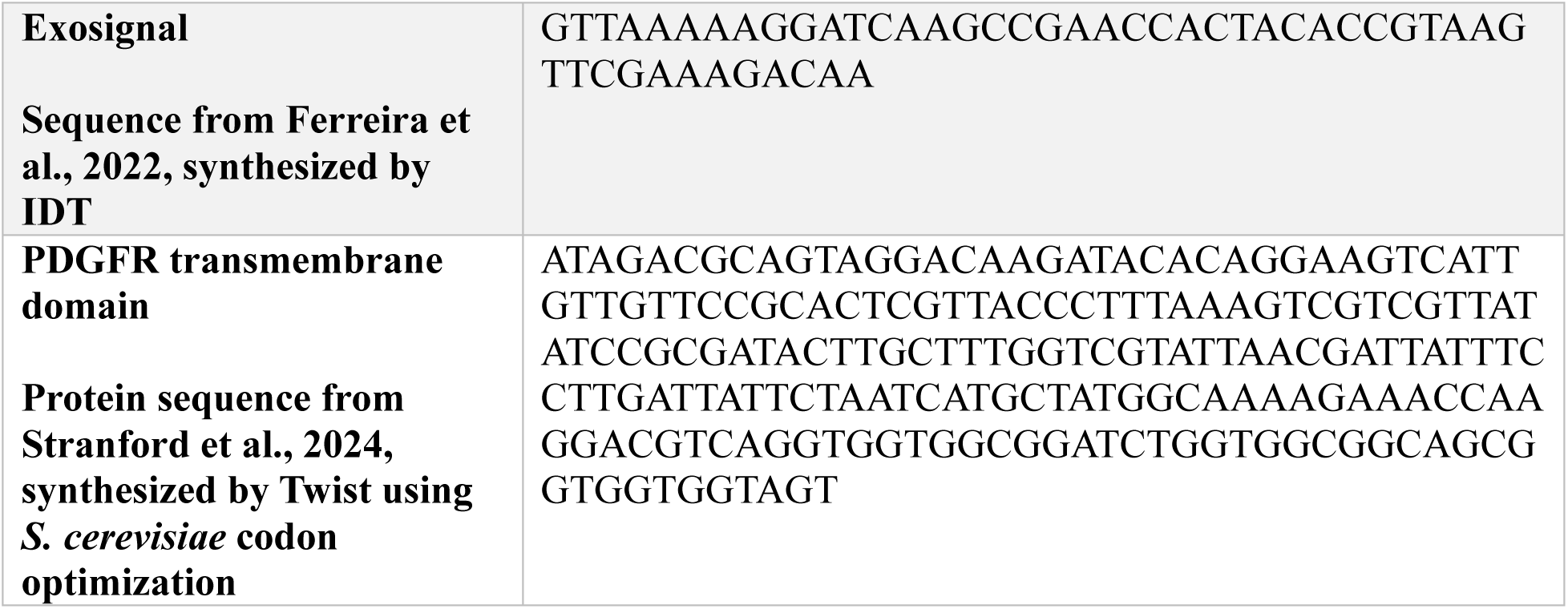
EVclo scaffold sequences used in this study. Asterisk after plasmid name indicates plasmids used to transform yeast

#### 2.1.1 | Sourcing of biological parts

All parts outside the EVclo CDSs were sourced from the Dueber Modular Cloning Yeast Toolkit (Addgene Plasmid Kit #1000000061) (Lee et al., 2015). All primers were ordered from IDT (Integrated DNA Technologies, Coralville, USA). Full-length *H. sapiens* codon CD63 was synthesized by IDT as a gBlock, flanked by Gateway attB sites, and combined with pDONR221 via Gateway BP cloning (described below) to generate a Gateway Entry clone plasmid.

Fragments of CD63 were PCR amplified from this plasmid, as described below. The *S. cerevisiae* codon-optimized PDGFR transmembrane domain was synthesized as a Gene Fragment from Twist (Twist Biosciences, Boston, USA), flanked by BsaI sites with part type 3b overhangs and BsmBI sites for insertion into the YTK part plasmid pYTK001 via Golden Gate cloning (described below). *H. sapiens* codon Exosignal polypeptide was ordered as 2 overlapping primers with part type 4a BsaI sites and BsmBI sites for insertion into pYTK001.

These fragments and part plasmids were then used in Golden Gate reactions as described below to create EVclo constructs. Bro1 was PCR amplified from the yeast genome as described below, inserted into the part plasmid pYTK001, and used from this plasmid to create EVclo constructs via Golden Gate cloning as below. The Gateway cloning cassette and EGFP were PCR amplified from plasmid pAG416GPD-ccdB-EGFP, a gift from Susan Lindquist (Addgene plasmid #14196) (Alberti et al., 2007), inserted into pYTK084 modified to serve as a part plasmid, and used from this plasmid to create EVclo constructs via Golden Gate cloning as below. The BbsI site introduced to the beginning of the type 3 and 3a Gateway cassettes was added using the PCR primer for insertion into pYTK001. Genomic integration used plasmids from the Markerless Yeast Localization and Overexpression (MyLO) CRISPR-Cas9 toolkit (Addgene Plasmid Kit #1000000210) (Bean et al., 2022). The MyLO integration vector pBBK96 was modified using primers to change ConLS to ConLS’ and ConR1 to ConRE’, to match the EVclo TU connectors that had been previously established, followed by Golden Gate cloning as below.

#### 2.1.2 | Polymerase Chain Reaction (PCR) amplification of parts

Yeast cell genomic DNA for PCR template was generated by suspending a yeast colony in 100 µL of 200 mM LiOAc with 1 % SDS, in a 1.5 mL Eppendorf tube. This tube was incubated on a heat block at 70°C for 15 minutes, and then 300 µL of 96% ethanol at room temperature was added. The tube was vortexed and centrifuged at 15,000 x g for 3 minutes. The supernatant was removed, the pellet was resuspended in 100 µL of 1X TE, and the tube was centrifuged at 15,000 x g for 1 minute. This supernatant was removed and transferred to a fresh 1.5 mL Eppendorf tube as yeast genomic DNA. If not used immediately, the yeast genomic DNA was stored at -20°C and thawed before use.

Plasmids for PCR templates were miniprepped using a QIAprep Spin Miniprep Kit (Qiagen, Cat#27106, Toronto, Canada), or an E.Z.N.A. Plasmid DNA Mini Kit I (Omega Bio-Tek, Cat#D6943, Norcross, USA), following manufacturer instructions. Plasmid purity and concentration was assessed using a NanoDrop 2000c spectrophotometer (Thermo Scientific, Wilmington, USA), and plasmid preparations were diluted to 1-10 ng/µL in MilliQ water for use as PCR template. If not used immediately, plasmids were stored at -20°C and thawed before use.

PCR reactions used 2X Phusion High-Fidelity PCR Master Mix with HF Buffer (ThermoFisher Scientific, Cat#F531, Waltham, USA). 20 µL reactions were created following manufacturer instructions, omitting DMSO, with 1 µL of each primer at 10 µM and 1 µL of yeast genomic or plasmid DNA as described above. Thermocycle parameters followed manufacturer instructions for a 3-step protocol, using the longest time indicated, with annealing temperatures between 50°C and 60°C.

PCR amplicons were assessed via DNA gel electrophoresis using 1-2% agarose gels in TAE and SYBRSafe DNA stain (ThermoFisher, Cat# S33102), with running parameters varying based on the size of the amplicon. Correct-sized bands were excised and purified using a QIAquick Gel Extraction Kit (Qiagen, Cat#28706), or an E.Z.N.A. Gel Extraction Kit (Omega Bio-Tek, Cat#D2500), following manufacturer instructions.

#### 2.1.3 | Golden Gate reactions

Golden Gate reactions were created at 10 µL volumes, using 1 µL of Fast Digest BpiI/BbsI (ThermoFisher, Cat#FD10114), Eco31I/BsaI (ThermoFisher, Cat# FD0294), or Esp3I/BsmBI (ThermoFisher, Cat#FD0454), 1 µL of 10x Fast Digest buffer supplied with the enzyme, 1 µL of T7 DNA ligase (New England Biolabs, Cat#M0318, Ipswich, USA), and 1 µL of 10 mM ATP (New England Biolabs, Cat#P0756). Plasmids were prepared via miniprep as described above, using part plasmids at 1 µL of each plasmid prep at 75 ng/µL concentration, or at 20 fmol of each part after restriction digest. PCR amplicons used in assemblies were gel purified as described above, and the eluted gel purification was used to bring the final reaction volume to 10 µL. If the final reaction volume was less than 10 µL, MilliQ water was added to bring the final reaction volume to 10 µL. Reactions used the following thermocycle parameters: Step 1: 37°C for 1 min, Step 2: 16°C for 1 min, Step 3: return to Step 1 30 times, Step 4: 37°C for 15 min, Step 5: 85°C for 15 min, Step 6: 4°C hold infinity. This thermocycler protocol is for end on digestion, if an end on ligation protocol was used Step 4 was replaced with 16°C for 60 min.

#### 2.1.4 | Gateway cloning reactions

Gateway cloning reactions were created at 10 µL volumes, using 1 µL of BP Clonase II enzyme mix (ThermoFisher, Cat#11789020), or 1 µL of LR Clonase II enzyme mix (ThermoFisher, Cat#11791020). For BP reactions, 1 µL of 150 ng/µL Donor vector was added. AttB flanked fragments for BP reactions produced by synthesis or PCR amplification had variable concentrations; 150 ng, or a maximum of 8 µL, of the fragment was used. For LR reactions, 1 µL of 150 ng/µL was used for both Destination vector and Entry clone. For both BP and LR reactions, if the final reaction volume was less than 10 µL TE buffer was added to bring the final volume to 10 µL. Reactions were incubated on the benchtop at room temperature overnight. 1 µL of 2 µg/µL proteinase K, provided with the Clonase enzyme mix, was then added, and the reaction was incubated at 37°C for 10 minutes.

#### 2.1.5 | Bacterial transformation

Golden Gate and Gateway reactions were transformed into DH5α *E. coli* bacteria, either made chemically competent in house, or bought from New England Biolabs (Cat#C2987H). For plasmids containing the Gateway cassette, ccdB-resistant TOP10 *E. coli* bacteria were used, either made chemically competent in house or bought from ThermoFisher (Cat#A10460). In all cases the same bacterial transformation protocol was used, following the New England Biolabs manufacturer instructions.

#### 2.1.6 | Validating plasmids

Single bacterial colonies were inoculated in 5 mL LB liquid with appropriate antibiotic selection and grown at 37°C overnight in a shaking incubator at 250 rpm. The next day, 1 mL of the overnight culture was reserved to plate and stock successful cultures, and the rest was miniprepped as described above. Miniprepped plasmids were analyzed by restriction digests followed by gel electrophoresis, and plasmids that looked correct were sent for whole plasmid sequencing performed by Plasmidsaurus using Oxford Nanopore Technology with custom analysis and annotation (Plasmidsaurus, Arcadia, USA).

### 2.2 | Yeast strains and reagents

*Saccharomyces cerevisiae* strains used are listed in Table 4. GFP knock–in clones are from a complete genome–wide strain collection purchased from Invitrogen Corp. (Cat# 95702, Carlsbad, USA; see Huh et al., 2003). Biochemical and yeast growth reagents were purchased from Sigma-Aldrich (Oakville, Canada), BioShop Canada Inc. (Burlington, Canada) or Thermo-Fisher Scientific (Burlington, Canada).

**Table 4 –.** Yeast strains used in this study.

| Strain | Genotype | Source |
| --- | --- | --- |
| **BY4741** | *MATα his3-Δ1 leu2-Δ0 met15-Δ0 ura3-Δ0* | Huh et al. |
| **CBY309** | BY4741, [*pTDH3-Exosignal-EGFP-tADH1 URA3*] | This study |
| **CBY311** | BY4741, [*pTDH3-mRuby2-PDGFR-EGFP-tADH1 URA3*] | This study |
| **CBY313** | BY4741, [*pTDH3-CD63(TM1-3)-mRuby2-CD63(TM4)-EGFP-tADH1 URA3*] | This study |
| **CBY326** | BY4741, *SSA2-GFP::His3MX FF18::pCCW12-NHX1-mRuby2-tADH1* | This study |
| **CBY327** | BY4741, *TOM70-GFP::His3MX FF18::pCCW12-NHX1-mRuby2-tADH1* | This study |
| **CBY330** | BY4741, [*pTDH3-Bro1-EGFP-tADH1 URA3*]<br>*FF18::pCCW12-NHX1-mRuby2-tADH1* | This study |
| **CBY332** | BY4741, [*pTDH3-Exosignal-EGFP-tADH1 URA3*]<br>*FF18::pCCW12-NHX1-mRuby2-tADH1* | This study |
| **CBY350** | BY4741, *FF18::pCCW12-NHX1-mRuby2-tADH1* | This study |
| **CBY351** | BY4741, [*pTDH3-Bro1-EGFP-tADH1 URA3*] | This study |
| **CBY352** | BY4741, *HIS3::pTDH3-Bro1-EGFP-tADH1* | This study |
| **CBY353** | BY4741, *HIS3::pTDH3-CD63(TM1-3)-mRuby2-CD63(TM4)-EGFP-tADH1* | This study |
| **Ssa2-GFP** | BY4741, *SSA2-GFP::His3MX* | Huh et al. |
| **Tom70-GFP** | BY4741, *TOM70-GFP::His3MX* | Huh et al. |

Unless otherwise stated, yeast cultures were grown at 30°C and 200 rpm in a shaking incubator (Infors HT Multitron II, Bottmingen, Germany) or on a shaker (Infors HT Orbitron) in a climate-controlled room. Unless otherwise stated, PBS used for washing cells was made in the lab, and 0.1 µm-filtered, tissue culture grade PBS (Cytiva, Cat#SH30028.02, Vancouver, Canada) was used for final resuspension for analyses. Unless otherwise stated, 50- and 15-mL Falcon tubes were centrifuged using an Eppendorf 5804R (Eppendorf, Enfield, USA) or an Allegra X-12R (Beckman Coulter, Mississauga, Canada), and 1.5 mL Eppendorf tubes were centrifuged using an Eppendorf 5424.

#### 2.2.1 | Genetic engineering of yeast strains

For all transformations, plasmids were prepared by miniprep as described above. For plasmid-based expression, 1 µg of plasmid DNA was used for yeast transformation. For genomic integration, 750 ng of donor plasmid was used, digested by NotI-HF (New England Biolabs, Cat#R3189L) restriction enzyme in a 10 µL reaction. For genomic integration, 500 ng of a MyLO Cas9 guide plasmid with split Kanamycin marker was used, digested by AatII (New England Biolabs, Cat#R0117S) and SbfI-HF (New England Biolabs, Cat# R3642S) restriction enzymes in a 10 µL reaction. Both donor and guide plasmids were digested either the night before or the morning of use, in PCR tubes in a thermocycler, with a protocol of 37°C for 1 hour, 80°C for 20 minutes, 80°C to 23°C at a ramp of 0.1°C/sec, followed by hold for infinity at 4°C. For EVclo TUs, the MyLO splitKanR guide plasmid pBBK17, targeting the HIS3 genomic locus, was used. For Nhx1-mRuby2 integration, the MyLO splitKanR guide plasmid pBBK11, targeting the FF18 genomic locus, was used.

Yeast colonies were inoculated into test tubes with 5 mL YPD or SD -URA selection media and grown overnight. Cultures were then back diluted to 0.4 OD_600nm_/mL in 10 mL YPD or SD - URA media in a 125 mL Erlenmeyer flask and grown for 4-6 hours until the OD_600nm_/mL reached 1.2-2.0. 3 OD units per transformation were transferred to 15 mL Falcon tubes and pelleted by centrifugation at 2,500 x g for 3 minutes, followed by removal of supernatant and resuspension of the cell pellet in 1 mL 1X TE buffer. Cell suspensions were transferred to 1.5 mL Eppendorf tubes and washed with 1X TE buffer by centrifugation at 4,500 x g – 10,000 x g for 1 minute. The washed cell pellet was resuspended in 1 mL 0.1M LiOAc and incubated on the bench at room temperature for 1 hour. Cells were pelleted by centrifugation at 4,500 x g – 10,000 x g for 1 minute, and the supernatant removed.

Transformation reagents were then added to the cell pellet, starting with 10 µL salmon sperm DNA (ThermoFisher, Cat# 15632011) that was boiled, frozen as an aliquot, and freshly thawed. Plasmid DNA was added next; for plasmid-based expression 1 µg of DNA was used, for genomic integration the full 10 µL restriction digests of donor and guide plasmids, without further purification, were used. MilliQ water was then added to make the final volume of plasmids and water 84 µL. 36 µL of 1 M LiOAc was added, followed by 240 µL PEG/PLATE solution (1 part 10X TE buffer, 1 part 1 M LiOAc, and 8 parts 50 % PEG3350). For some transformations, 36 µL DMSO was also added. Transformation tubes were vortexed for 2 seconds and incubated on the bench at room temperature for 30 minutes followed by incubation in a 42°C water bath for 30 minutes, with 2 second vortexing every 15 minutes. Finally, the tube was incubated at room temperature on the bench for 5 minutes.

Cells were pelleted at 10,000 x g for 1 minute. For plasmid-based expression, cell pellets were resuspended in 100-150 µL MilliQ water or growth media, several dilutions were plated on SD - URA selection plates using 4.5 mm glass beads, and plates were incubated stationary at 30°C. For genomic integration, cell pellets were resuspended in 500 µL YPD or SD -URA selection media and incubated stationary at 30°C overnight. The next day, 150 µL of the culture, and several dilutions, were plated on SC +G418 or SD -URA +G418 solid media, both made with Yeast Nitrogen Base without ammonium sulfate, supplemented with L-glutamic acid monosodium salt hydrate, for strong G418 selection. For both types of transformation, plates were checked 2 and 3 days after plating for colonies. Colonies on the plate were screened on an Invitrogen Safe Imager 2.0 Blue-Light Transilluminator (ThermoFisher) for GFP expression.

Positive colonies were inoculated in 5 mL SC or SD -URA liquid media and cultured overnight in test tubes. Cultures were screened for GFP or RFP using a widefield microscope (Leica DMI6000B, Wetzlar, Germany), and a plate reader (BioTEK Synergy H1, Winooski, USA) in some cases, and positive cultures were used as templates for PCR using primers for the plasmid or genomic locus. The brightest positive cultures were selected and streaked on plates and cryogenically stocked for future experiments.

### 2.3 | Yeast cell assays

#### 2.3.1 | Yeast cell microscopy

Yeast colonies were inoculated in 5 mL SC or SD -URA in test tubes and grown overnight. Cultures were then back diluted by transferring 500 µL of culture to 5 mL of fresh media in test tubes. After growing for 3.5-4 hours, cultures were measured for OD_600nm_/mL, and 5 OD units of culture were transferred to 1.5 mL Eppendorf tubes. Cultures were washed 2x in 1 mL PBS by centrifugation at 10,000 x g for 1 min, with final resuspension in 100 µL PBS (Cytiva). 7 µL of cell suspension was transferred to a 15 mm x 25 mm 1-2 % agarose pad of lab tape thickness on a microscope slide and covered with a 24 mm x 60 mm #1.5 glass coverslip. Cells were imaged on a Zeiss (Carl Zeiss, Toronto, Canada) Axio confocal microscope with a CICERO (Crest Optics, Rome, Italy) spinning disk, using a 100x Plan Apo NA 1.46 oil immersion objective.

Images were acquired using a Hamamatsu (Hamamatsu, Japan) Flash 4.0 LT Plus sCMOS camera, model C11440-42U30, with 16-bit depth, using the full chip at 2048 x 2048 pixels. Brightfield images were acquired using white transmitted light with a 100 ms exposure time. For fluorescence excitations, an 89 North (Williston, USA) LDI-5 laser launch was used, with a 488 nm laser exciting the GFP channel and a 555 nm laser exciting the RFP channel. Fluorescence emissions were collected using filters; 525/50 for GFP and 600/50 for RFP. Most strains were imaged at 60 % laser power and 400 ms exposure time for the GFP channel and 100 % laser power and 800 ms exposure time for the RFP channel, except the mRuby-PDGFR-GFP strain that was imaged at 100 % laser power and 2800 ms exposure time in the RFP channel, and Tom70-GFP that was imaged at 100 % laser power and 1200 ms exposure time in the GFP channel. Volocity software, version 7.0.0, was used to control image acquisitions.

#### 2.3.2 | Yeast cell flow cytometry

Yeast colonies were inoculated in 5 mL SC or SD-URA in test tubes and grown overnight. Cultures were then back diluted by transferring 500 µL of overnight culture to 5 mL of fresh media in test tubes. After growing for 3.5-4 hours, cultures were measured for OD_600nm_/mL, and 5 OD units of culture were transferred to 1.5 mL Eppendorf tubes. Cultures were washed 2x in 1 mL PBS by centrifugation at 10,000 x g for 1 min, with final resuspension in 1 mL PBS (Cytiva). 250 µL of suspension was added to 750 µL of PBS (Cytiva) in a deep well 96 well plate and transferred to an Accuri C6 flow cytometer with an autosampler (BD Biosciences, Mississauga, Canada), controlled by BD Accuri C6 software version 1.0.264.21. Wells were sampled using the slow fluidics setting, with a flow rate of 14 µL/min and a core size of 10 µm, with 1 wash cycle between each well, and an event threshold of 80,000 on the FSC-H channel. 100,000 events were collected for each sample. GFP fluorescence signals for events were captured using the FL1 channel, with excitation using a 488 nm laser and emissions collected using a 533/30 filter.

Yeast cell flow cytometry data was analyzed using the Floreada.io website, last updated 06 August 2024, accessed in November and December 2024. Flow cytometry data analysis started by gating out debris by examining scatter plots with FSC-A as the x-axis and SSC-A as the y-axis and drawing a gate to exclude outliers. A singlet gate was then drawn by examining scatter plots with FSC-A as the x-axis and FSC-H as the y-axis and drawing the gate to exclude points not on the diagonal of the main body of the data points. Finally, a GFP+ gate was set by examining the WT(BY4741) singlet data histogram with FL1-H as the x-axis, arriving at 1x10^3^ FL1-H as the GFP+ cutoff used for all data. The percent GFP+ events and median FL1-A intensities of the GFP+ gate were exported from the Floreada.io website as a CSV file and loaded into Microsoft Excel for further processing. Samples were assayed in duplicate and averaged to provide the value for that biological replicate.

#### 2.3.3 | Yeast cell colocalization analysis

Yeast colonies were inoculated in 5 mL SC media in test tubes and grown for 8 hours. The OD_600nm_/mL for each culture was measured, and cultures were inoculated in 30 mL of SC media in 125 mL Erlenmeyer flasks using the formula: Volume = (0.5^(17/1.55))*(6/OD_600nm_)*30, multiplied by an empirically determined, strain-specific factor between 2 and 3.5. Cultures were grown for 17 hours and measured for OD_600nm_/mL. 100 OD units of culture were transferred to a 15- or 50-mL Falcon tube and washed 2x with 1 mL PBS by centrifuging at 2,830 x g for 3 minutes, with a final resuspension in 1 mL PBS (Cytiva). 250 µL of suspension was transferred to each of (2) 1.5 mL Eppendorf tubes. Prior to imaging, cells were pelleted by centrifugation at 2,850 x g for 3 minutes and cell pellets were incubated on the bench at room temperature for 15 minutes. The pellets were then resuspended in 100 µL PBS (Cytiva) and incubated on the bench at room temperature for 15 minutes. 5 µL of suspension was then transferred to a 15 mm x 25 mm 1-2 % agarose pad of lab tape thickness on a microscope slide and covered with a 24 mm x 60 mm #1.5 glass coverslip. Cells were imaged on a Zeiss Axio confocal microscope with a CICERO spinning disk, as described above.

To analyze images for colocalization, regions of interest (ROI) outlining individual cells were obtained using the YeastSpotter neural network (Lu et al., 2019) or were manually drawn. Prior to analysis, fluorescent channel images were background subtracted using the built-in Fiji (ImageJ 1.54f) function with a 50-pixel rolling ball radius. Colocalization analysis was performed using the Fiji BIOP JACoP plugin (Bolte & Cordelières, 2006), with the thresholded Manders’ colocalization coefficient (tM) as a readout. The tM measures how much of the mRuby fluorescence intensity above threshold in that ROI (Ʃ mRuby) is found in pixels that are also above the threshold for GFP (Ʃ mRuby, coloc). To determine the thresholds and establish a lower bound for the tM, we analyzed the background and noise in micrographs for each strain by moving the cell ROIs to regions of the image without cells. The mean pixel intensity of these blank ROIs was calculated in both GFP and mRuby channels for each ROI, and the thresholds for the BIOP JACoP plugin were calculated as the mean of all blank ROIs plus 3 times the standard deviation over all blank ROIs for each channel. Using these thresholds, the blank and cell ROIs were then analyzed to calculate tM. The BIOP JACoP plugin provides a p-value to determine the significance of colocalization using Costes’ randomization, here we used a block size of 5 and 100 randomizations. ROIs showing a Costes’ correlation p-value less than 1 were discarded from further analysis. Results were exported from Fiji as a CSV file and transferred to Microsoft Excel for additional processing and analysis. The mean pixel intensities of all cell ROIs, after background subtraction, in both GFP and RFP channels, were also calculated and collected in a CSV output file, followed by subtraction of the colocalization analysis threshold value using Microsoft Excel.

#### 2.3.4 | Yeast whole cell lysate for Western blot analysis

Whole cell lysates were prepared by collecting 1 OD_600nm_ unit of yeast cells from 8-hour seed cultures remaining after inoculation for EV isolation for Western blot analysis (see below).

Culture was transferred to 1.5 mL Eppendorf tubes and cells were pelleted by centrifugation at 3,500 x g for 5 minutes at room temperature. Pellets were resuspended in 10 % trichloroacetic acid (TCA) at 4°C and incubated on ice for 1 hour. Samples were then centrifuged at 12,000 x g for 5 minutes and pellets were washed with 0.1 % TCA at 4°C and resuspended in 100 µL boiling buffer (1.5 M Tris pH 8.5, 0.5 M EDTA, 10 % SDS) with 1 µL protease inhibitor cocktail (Abcam, Cat# ab271306, Waltham, USA). 0.5 mm glass beads (BioSpec Products, Cat#11079105, Bartlesville, USA) were added to the 0.1 mL mark on the centrifuge tube, and samples were subjected to 5 minutes in a cell disruptor (Disruptor Genie, Scientific Industries) at 4°C in a climate-controlled room, followed by incubation at 95°C in a heat block for 5 minutes. 100 µL of urea buffer (150mM Tris pH 6.8, 6M Urea, 6 % SDS, 40 % Glycerol, 100mM DTT, 0.01 % Bromophenol blue) with 1 µL protease inhibitor cocktail was added, samples were disrupted again for 5 minutes at 4°C and incubated at 95°C for 5 minutes. Samples were stored at 4°C prior to analysis by SDS-PAGE, with 15 µL of sample used for Western blot analysis as described below.

### 2.4 | EV isolation, staining, and purification

#### 2.4.1 | Ultrafiltration

EVs were isolated using differential centrifugation followed by ultrafiltration using Amicon Ultra-15 centrifugal filters with a 100 kDa MWCO (Millipore, Cat#910096, Tullagreen, Ireland). Filter units were reused up to 3 times within 1 month and stored filled with PBS (Cytiva) at 4°C between uses. Yeast liquid seed cultures were prepared by inoculating colonies into 15 mL of YPD medium in 125 mL Erlenmeyer flasks and grown for 8 hours. Seed cultures were measured for OD_600nm_/mL and used to inoculate 1 L of YPD media in 2 L Erlenmeyer flasks using the formula Volume = (0.5^(17/1.55))*(6/OD_600nm_)*1000, multiplied by an empirically determined, strain-specific factor between 2 and 3.5. Cultures were grown for 17 hours, or until cultures reached densities of 6 – 10 OD_600nm_/mL. Cultures were then transferred to 500 mL PPCO centrifuge bottles (Nalgene, ThermoFisher Scientific, Cat# 21020-050) and pelleted by centrifugation (Avanti J26XPI or Avanti J30I centrifuge, JA-10 rotor, Beckman Coulter) at 3,000 x g, max acceleration and deceleration, for 10 minutes at room temperature. Supernatants containing EVs indiscriminately released during growth were discarded, and cell pellets were washed twice with PBS. Cell pellets were then exposed to heat conditioning at 42°C for 15 minutes in a water bath, immediately resuspended in 25 mL of PBS (Cytiva), and incubated for an additional 15 minutes at 42°C. Concentrated yeast samples in PBS were transferred to 50 mL polycarbonate centrifuge bottles (Beckman Coulter, Cat# 357002) that were pre-chilled on ice. All subsequent steps were carried out on ice or at 4°C. Resuspended cell pellets were centrifuged (Avanti J26XPI or Avanti J30I centrifuge, JA-25-50 or JA-30.50 Ti rotor, Beckman Coulter) at 15,000 x g, max acceleration and deceleration, for 15 minutes. The resulting supernatant was recovered and filtered using pre–chilled 0.22 µm filters (FroggaBio, Cat#SF0.22PES, Toronto, Canada) to remove cell debris. 15 mL of this filtered supernatant was transferred to an Amicon filter, and centrifuged (Eppendorf 5804R or Allegra X-12R) at 2,500 x g for 15 minutes at 4°C. The flowthrough was discarded and the remaining filtered supernatant was transferred to the filter and centrifuged using the same parameters for an additional 10 minutes. If the remaining volume in the top of the filter unit was > 200 µL additional 1 minute centrifuge steps were used to bring the volume below 200 µL. Concentrated EV samples were recovered from the top of the Amicon filter unit by pipetting the sample over the bottom and lower sides of the filter using a sweeping motion. EV samples were kept on ice or at 4°C prior to analysis or transferred to 4°C for storage.

#### 2.4.2 | FM4-64 staining and size exclusion chromatography

Some EV samples were stained with FM4-64 to label EVs, using 2 µL of 2.5 mM FM4-64 added to 150 µL of EV sample in a brown 1.5 mL Eppendorf tube and incubated at 37°C for 15 minutes, followed by size exclusion chromatography purification using Izon Gen2 qEV single 70 nm columns (Izon, Cat#ICS70P-1621, Christchurch, New Zealand). Columns were reused up to 5 times within three months, with cleaning, storage, and preparation for use according to manufacturer instructions. For use, 130-150 µL of EV sample was added to the column, allowed to pass the frit, and then 850-870 µL of PBS (Cytiva) was added to the column. When this volume passed the frit, 800 µL of PBS was added, and the flowthrough was collected in a 1.5 mL Eppendorf tube as the purified EV sample.

### 2.5 | EV characterisation

#### 2.5.1 | EV particle size and concentration measurements

To measure particle size and concentration, we conducted nanoparticle tracking analysis (NTA) using a ZetaView PMX120 nanoparticle tracking analysis instrument (Particle Metrix, Ammersee, Germany) with software version 8.0.5.16 SP7, or a PMX130 instrument with software version 8.06.01. EV samples were diluted between 1:25 to 1:2000 in PBS (Cytiva). 1 mL of diluted EV samples was manually loaded with a 1 mL syringe and samples were slowly injected until conditions appeared optimal for acquisition. Both ZetaView instruments were used with the same settings: Temperature (23°C), laser λ (488 nm), filter λ (scatter) sensitivity (75), shutter (100), frame rate (30 fps), cycles (2), positions (11), and minimum trace length (30). Unless otherwise reported, EV samples were measured twice to provide 2 technical replicates for each biological replicate, and samples from ≥ 3 different yeast cultures (biological replicates) were examined.

### 2.5.2 | EV particle yield calculation

For each flask of yeast culture used for EV isolations, the OD_600nm_/mL was measured immediately prior to isolation. Cell numbers were calculated by multiplying culture OD_600nm_ /mL by 3.6 x 10^7^ cells/OD_600nm_ unit. The EV sample particle concentration in particles/mL was obtained from NTA as above and multiplied by the EV sample volume to give the total number of particles recovered. The total number of particles in the EV sample was divided by the total number of cells to determine number of particles released per cell.

#### 2.5.3 | EV transmission electron microscopy

EV samples were prepared for TEM on Formvar/Carbon 300 mesh copper grids (Electron Microscopy Science, Cat# FCF300-Cu-50). Fixative was prepared fresh before sample prep by adding 5 µL 50% glutaraldehyde (photographic grade) to 95 µL 0.1 M sodium cacodylate. 5 µL of EV sample was pipetted onto a grid and allowed to adsorb for 5 minutes at room temperature. 5 µL of glutaraldehyde/sodium cacodylate fixative was then added to the EV sample drop and left at room temperature for 5 minutes. 5 µL of 0.15 M glycine was added to the drop, left for 1 minute, and the drop was wicked away using the edge of a folded kimwipe. 5 µL of 0.15 M glycine was then added to the grid and left for 2 minutes, followed by 10 µL of MilliQ water left for 1 minute and then wicked away. Another 10 µL of MilliQ water was added to the grid, left for 1 minute, and then wicked dry. Finally, 5 µL of 1 % phosphotungstic acid in MilliQ water was added to the grid as a negative stain, left for strictly 30 seconds, and wicked dry. Grids were transferred to parafilm in a glass Petri dish, covered, and allowed to dry on the bench at room temperature overnight. Grids were imaged at 200 kV using a Talos F200X G2 (S)TEM (ThermoFisher Scientific, Toronto, Canada) with a Ceta 16M 4k x 4k CMOS camera, at 11,000 or 36,000x magnification.

#### 2.5.4 | EV fluorometry

EV fluorometry experiments assayed 3 strains for a given experiment, isolating EVs from 3 L of culture spun to a single pellet for each strain, using the ultrafiltration protocol above. EV sample concentration was measured using NTA as above, and the minimum number of particles across samples was determined. More concentrated EV samples were diluted so that all 3 samples had the same number of particles in 650 µL PBS (Cytiva). A black, V-well, 96-well plate (ThermoScientific Nunc Cat#249945) had every well except the first column filled with 150 µL of PBS (Cytiva). 300 µL of EV sample or 2.5 nM fluorescein in PBS was added to the first column wells, and (10) 1:2 serial dilutions moved 150 µL of concentrated sample to the next well and mixed, leaving the last column as blanks. Each EV sample and the fluorescein curve were run in duplicate, providing two measurements for each concentration. The plate was read using a BioTEK Synergy H1 plate reader, at room temperature, using an excitation wavelength of 485 nm, an emission wavelength of 515 nm, and a gain of 130. Results were exported as an Excel file and transferred to a second Excel file for processing and conversion to calibrated molecules of equivalent fluorescein (MEFL) units as described in Supplemental Figure 1 (Vignoni et al., 2019). The resulting values for the 2 technical replicates were averaged to generate a single value for each biological replicate.

#### 2.5.5 | EV Western blot analysis

EV samples for Western blot analysis were isolated from 2 L of culture spun to a single pellet for each strain, using the ultrafiltration followed by the size exclusion chromatography (SEC) protocols above. 8 µL of protease inhibitor cocktail (Abcam, Cat# ab271306) was added to the 800 µL sample retrieved from the SEC column, and the sample was stored at 4°C overnight.

Samples were then concentrated to roughly 15 µL using a 0.5 mL 10 kDa MWCO Amicon filter (Millipore, Cat# 501024). 5 µL of 4x Laemmli sample buffer (Bio-Rad, Cat#1610747, Mississauga, Canada) supplemented with protease inhibitor cocktail (Abcam, Cat# ab271306) and 50 mM DTT was added, and the sample was incubated at 95°C for 5 minutes prior to loading onto a gel.

Protein samples were separated by 10 % SDS-PAGE, transferred to PVDF membranes using a Bio-Rad Trans-Blot Turbo transfer system, and incubated with 5 % skim milk in 1x TBST on a nutator (Reliable Scientific) for 1 hour at room temperature. Membranes were then transferred to 5 % milk in 1x TBST containing anti-GFP mouse monoclonal antibody (MilliporeSigma, Cat#11814460001, Mannheim, Germany) at 1:1,000 dilution and incubated on a nutator at 4°C for 24 hours. Membranes were washed 5 times with 1x TBST, incubated for 45 minutes on a nutator at room temperature with 5 % milk in 1x TBST containing horseradish peroxidase-labeled affinity purified goat anti-mouse IgG antibody (SeraCare Cat#5450-0011, Milford, USA) at 1:10,000 dilution, and then washed an additional 5 times. Chemiluminescence of stained membranes was detected using a GE Amersham Imager 600 instrument (GE HealthCare, Piscataway, USA) and Amersham ECL Select Detection Reagent (Cytiva).

#### 2.5.5 | EV nano-flow cytometry

EV samples for nano-flow cytometry experiments were isolated from 2 L of culture spun to a single pellet for each strain, using the ultrafiltration followed by the FM4-64 staining and size exclusion chromatography protocols above. EV sample concentrations were measured using NTA as above. Samples were diluted to 3-5 x 10^7^ particles/mL in 500 µL PBS (Cytiva) and assayed on a CytoFLEX nano-flow cytometer (Beckman Coulter, Mississauga, Canada). EV samples were run for 10 minutes at a flow rate of 10 μL/min, with washes between samples. Acquisitions used gains of 200 for FSC, 100 for SSC, 275 for Violet SSC, 1000 for FITC, and 1000 for PC7 detectors, and a manual threshold of 1300 on the Violet SSC-H channel. After samples were assayed the remaining 400 µL of sample was recovered and 40 µL of 10% Triton X-100 was added. The sample was then vortexed and incubated on the benchtop at room temperature for at least 10 minutes and then examined on the CytoFLEX using the same parameters. Gates for fluorescence were established using CytExpert (Beckman Coulter, Version 2.5.0.77); the GFP positive gate was determined using WT(BY4741) EV sample, and the FM4-64 positive gate was determined using PBS +FM4-64 sample. Gates were transferred to the Floreada.io website (last updated 06 August 2024, accessed in November and December 2024) for further analysis and plotting. The number of events in GFP or FM4-64 positive gates were normalized to the number of total events between 1.5 x 10^3^ and 1 x 10^6^ on Violet SSC-H and 275 and 1075 on Violet SSC-Width.

### 2.6 | Data analysis and presentation

Confocal microscopy, transmission electron microscopy, and Western blot images were rotated, cropped, adjusted for brightness and contrast, inverted, annotated, and/or had color channels merged using Fiji/ImageJ version 1.54f. Western blot images were additionally adjusted for display using the Unsharp Mask Filter in Fiji. Calculations and compiling of data were carried out in Microsoft Excel version 2501. All graphs were plotted and analyzed for statistical significance using GraphPad Prism software version 10.4.1, with bars and statistical tests used described in figure legends. Values shown are mean ± S.E.M unless otherwise stated.

## 3 | RESULTS

### 3.1 | Optimizing synthetic biology-based cloning methods to modify yeast EVs

Taking a synthetic biology–based approach named “EVclo”, we initiated a proof–of–concept study to genetically engineer designer EVs in *S. cerevisiae* based on the Yeast Toolkit (YTK) for modular cloning (MoClo; Lee et al., 2015). The YTK assembly standard defines 11 part types (e.g. all promoters are type 2 parts) each flanked by unique four base pair overhangs generated when part plasmids are cut using BsaI, a type IIS restriction enzyme. Overhangs of each part type interface with adjacent part types, enabling directed assembly of expression plasmids using one-pot Golden Gate reactions. To generate transcriptional units (TUs) for each EV scaffold tested (see Figure 1a), we created part plasmids encoding the EV scaffold gene or “cargo” genes (mRuby2, EGFP) to be fused at the 5’or 3’ end of the scaffold gene. These multi-part coding DNA sequences (CDS) represent a single translational fusion. All other genetic modules were sourced from YTK part plasmids, including a strong constitutive promoter (pTDH3) and tADH1 terminator (Figure 1a top). The promoter, CDS, and terminator form all TUs to be expressed by yeast cells. TUs were flanked by YTK connector parts (ConLS’, ConRE’) allowing them to be moved between vectors using BsmBI Golden Gate reactions.

For plasmid-based expression (Figure 1a), TUs were inserted into backbones containing the high copy number yeast origin of replication (2µ) and URA3 module for auxotrophic selection. For genomic integration, we used parts from the Markerless Yeast Localization and Overexpression (MyLO) toolkit (Bean et al., 2022) featuring ∼125 bp homology arms for 7 different landing pad loci in the yeast genome. Both backbones feature ConLS’ and ConRE’ connector parts to enable transfer of TUs, as well as the ColE1 bacterial origin of replication and AmpR ampicillin resistance module to propagate plasmids in *E.coli*.

We hypothesized that engineered yeast cells will produce designer EVs following the schematic in Figure 1b: Transformation for plasmid-based expression (top) used media lacking uracil for selection. Transformation for genomic integration (bottom) used a second MyLO plasmid coding for Cas9, the guide sequence targeting the HIS3 locus, and a split kanamycin resistance module. Plating on media containing G418 selected for successful recombination, followed by screening of colonies for successful genomic integration. Yeast cells with TUs will express EV scaffold proteins fused to fluorescent proteins (cargoes). Successful scaffolds will be sorted into intralumenal vesicles (ILVs) within the multivesicular body (MVB), a site of EV biogenesis.

When the MVB fuses with the plasma membrane, ILVs containing scaffold fused to cargoes are released as designer EVs.

To replace fluorescent proteins with bioactive cargo proteins for future applications, we introduced Gateway cloning cassettes into our EVclo system (Figure 1c). This enables creation of Gateway Expression plasmid libraries by reacting a common Destination Vector with collections of Gateway Entry Clones containing different cargo genes via Gateway LR cloning. Gateway cloning cassettes (Figure 1c top) are composed of attR1 and attR2 recombination sites, flanking a CamR chloramphenicol resistance module for selection and a ccdB bacterial toxin module to increase the efficiency of Gateway cloning reactions. When combined with Gateway Entry clones (Figure 1c bottom) featuring a cargo gene part flanked by attL1 and attL2, reciprocal recombination sites of a Gateway LR reaction, the att sites recombine and the cargo part replaces the CamR and ccdB modules, in frame and fused with a flanking scaffold. To introduce a fourth CDS part, we incorporated a BbsI recognition site at the 5’ end of the type 3 and type 3a Gateway cloning cassettes. All constructs in this study featured test cargoes EGFP or mRuby2 inserted into the CDS using Gateway LR cloning.

To test our EVclo system, we selected Bro1 – an ALIX ortholog in *S. cerevisiae* – as a candidate scaffold because it was detected in yeast EVs (Logan et al., 2024). Given that machinery implicated in small EV biogenesis is conserved (i.e. ESCRTs), we also tested human EV scaffolds commonly used by other researchers. These include CD63 (Stickney et al., 2016), the PDGFR transmembrane domain (Stranford et al., 2024), and the small polypeptide ExoSignal (Ferreira et al., 2022) (Figure 1d). To fully validate this system, we designed and generated CDSs using a variety of part types, and tested CDSs with 2, 3 or 4 components that cover multiple strategies for EV cargo protein loading.

We tested two 2–component CDSs, the first containing the full length Bro1 as a type 3 part and the Gateway cloning cassette as a type 4a part, with EGFP Gateway cloned in to generate a C-terminal fusion. The Bro1-GFP TU was expected to generate a lumenal EV protein, and was tested using both plasmid-based expression, Bro1-GFP (P), and genomic integration, Bro1-GFP (I). The second contains the Gateway cloning cassette as a type 3 part and the *H. sapiens* codon ExoSignal as a type 4a part, with EGFP Gateway cloned in to generate an N-terminal fusion. The GFP-ExoSignal TU was expected to generate a lumenal EV protein and was only tested with plasmid-based expression.

The 3–component CDS included the Gateway cloning cassette as a type 3a part, the *H. sapiens* PDGFR transmembrane domain and a GS linker as a type 3b part, and EGFP as a type 4a part, with mRuby2 Gateway cloned into the 3a N-terminal position. The mRuby-PDGFR-GFP TU was expected to generate a membrane localized EV protein with mRuby2 on the EV surface and EGFP in the lumen, and it was only tested with plasmid-based expression. For the 4–component CDS, the *H. sapiens* CD63 gene was split in two at a site within the extracellular loop between transmembrane domains 3 and 4 (see Stickney et al., 2016). It encodes the N-terminal portion of CD63 introduced into the BbsI site upstream of the Gateway cassette as a type 3a part, the C-terminal portion of CD63 as a type 3b part, and EGFP as a type 4a part. mRuby2 was Gateway cloned into the 3a position. The resulting TU, called CD63*-mRuby-GFP, was expected to produce a membrane localized EV protein with mRuby2 on the EV surface and EGFP in the lumen, and it was tested using both plasmid-based expression and genomic integration.

### 3.2 | Variable expression of CDSs encoding EV scaffolds in *S. cerevisiae*

With genetically engineered yeast strains in hand, confirmed by fluorescence microscopy (Figure 2a), we first examined expression levels of EGFP-tagged fusion proteins within the cell population by flow cytometry (Figure 2b – d). We focused on EGFP, and not mRuby2, because it was fused to all candidate scaffolds allowing assessment of relative expression.

**Figure 2.**
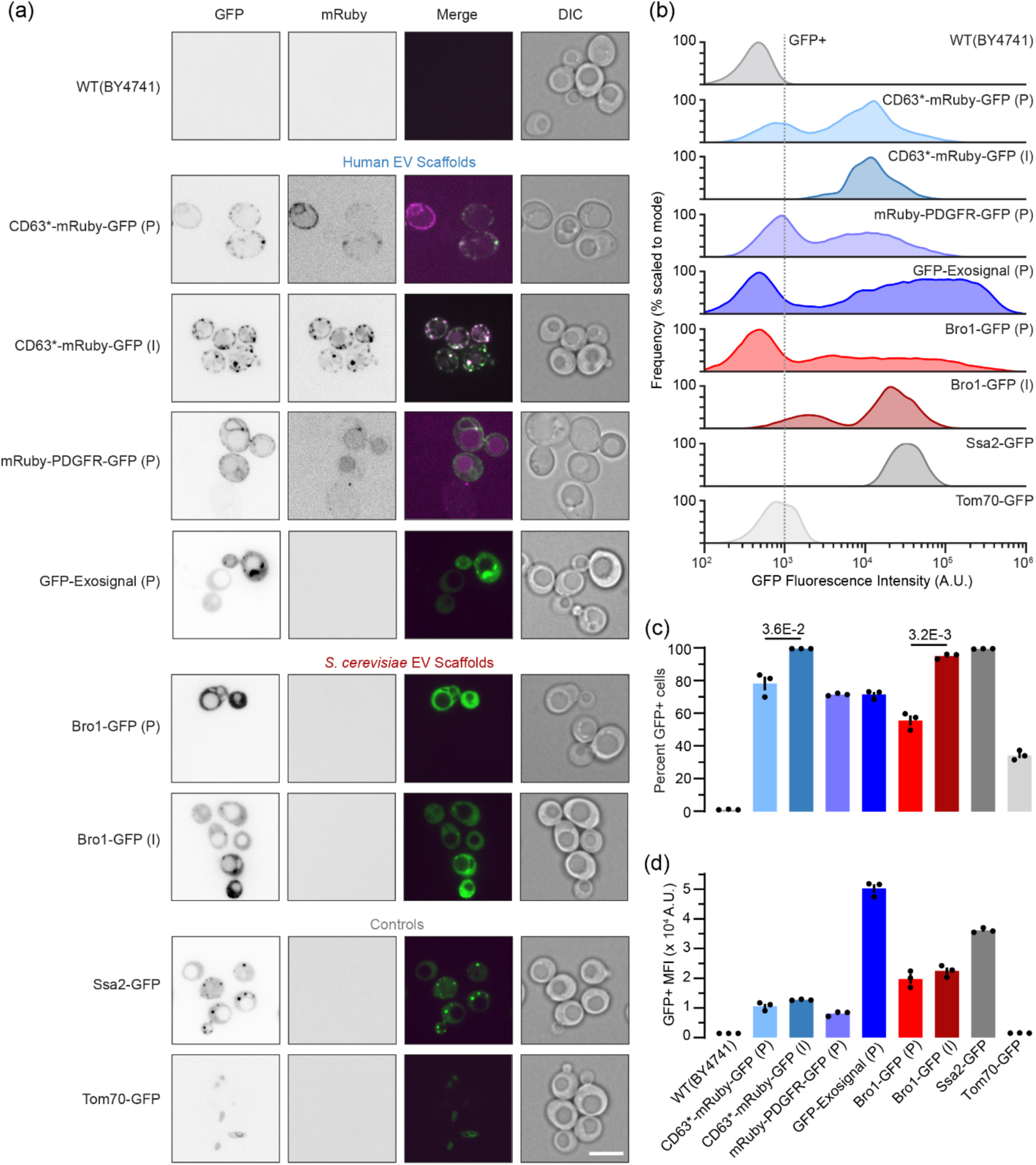
EVclo TUs express scaffold candidate proteins in *S. cerevisiae* as expected. (a) Representative confocal micrographs of yeast cells expressing EVclo constructs. Yeast cells for all strains were imaged in GFP, mRuby (RFP), and DIC channels. A merge of the GFP and mRuby channels is also presented. Strains with a (P) indicate plasmid expression, strains with an (I) indicate genomic integration. Ssa2-GFP, the HSP-70 ortholog previously demonstrated as an *S. cervisiae* EV cargo, tagged with GFP at the native genomic locus from the GFP library, serves as a positive control for EV experiments. Tom70-GFP, a mitochondrial protein not expected to appear in EVs, tagged with GFP at the native genomic locus from the GFP library, serves as a putative negative control for EV experiments. Scale bar, 5 µm. (b) Representative whole cell flow cytometry histograms of yeast strains tested. 1x10^6^ events were acquired, data presented are after gating for debris and singlets, normalized to the mode of the histogram. (c) Quantification of the percent GFP+ cells from flow cytometry data. After gating for debris and singlets, 1x10^3^ A.U. was set as the GFP+ threshold and the percentage of particles above this threshold was calculated. Points represent biological replicates, with the bar representing S.E.M. All strains show significant %GFP+ cells compared to WT(BY4741), with the highest p-value of 2.6E-3 for WT(BY4741) vs. Tom70-GFP via Welch’s 2-tailed t-test. Presented p-values compare plasmid expression vs. genomic integration via Welch’s 2-tailed t-test. (d) Median Fluorescence Intensity (MFI) of GFP+ populations from flow cytometry data. After gating for debris and singlets, the median fluorescence intensity of GFP+ events (>1x10^3^ A.U.) was calculated. Points represent biological replicates, with the bar representing S.E.M. All strains show significant GFP MFI compared to WT(BY4741), with the highest p-value of 1.1E-2 for WT(BY4741) vs. Tom70-GFP via Welch’s 1-tailed t-test.

Autofluorescence detected in wild type cells (without EGFP) was used to determine the threshold at which cellular EGFP fluorescence was reliably detected over background (> 1,000 A.U.). As controls, we also examined existing yeast strains harboring EGFP in their genome fused to either SSA2, an Hsp70 ortholog packaged within EVs (Logan et al., 2024), or TOM70, a mitochondrial outer membrane protein hypothesized to be excluded from EVs (Huh et al., 2003). We found that 99.7 ± 0.1 % of cells showed relatively high Ssa2-GFP fluorescence intensity, whereas cellular fluorescence intensity of Tom70-GFP was relatively low and detected in 34.5 ± 1.7 % of cells (Figure 2b and c). This observation is consistent with published data indicating that yeast cells contain ∼364,000 copies of Ssa2-GFP compared to 45,300 copies of Tom70-GFP (Huh et al., 2003), and thus Tom70-GFP+ cells are likely more difficult to detect by flow cytometry.

Despite use of a common strong, constitutive promoter, we found that the distribution of cellular fluorescence intensity observed with the population varied considerably depending on the CDS when encoded on a plasmid. ExoSignal-GFP showed the highest cellular fluorescence intensity and was detected in 71.8 ± 1.4 % of the yeast cell population (Figure 2c and d). Overall, Ssa2 = CD63 > ExoSignal > PDGFR > Bro1 > Tom70 in terms of percent GFP+ cells (Figure 2c), and ExoSignal > Ssa2 > Bro1 > CD63 > PDGFR > Tom70 for average cellular fluorescence intensity (Figure 2d). We found that ectopic expression of human proteins in *S. cerevisiae* did not impact translation, as human GFP-ExoSignal mean cellular fluorescence intensity exceeded *S. cerevisiae* Bro1-GFP when behind the same promotor. Lumenal EV fusion proteins showed higher average cellular fluorescence intensity compared to transmembrane proteins. Compared to plasmid expression, genomic integration of CDSs for CD63*mRuby-GFP and Bro1-GFP significantly increased the proportion of cells that expressed detectable GFP from 78.5 to 99.8 % and 55.8 to 95.3 %, respectively, but did not show a significant change in average cellular fluorescence intensity.

### 3.3 | Candidate scaffolds localize to predicted intracellular sites of EV biogenesis

We next visualized live cell populations using confocal fluorescence microscopy (Figure 3a). All genetically modified strains showed intracellular fluorescence over background (unmodified wild type cells) and relative intracellular fluorescence correlated with flow cytometry data, whereby brightest signals were observed in cells expressing soluble EV scaffolds GFP-ExoSignal or Bro1-GFP. In addition, intracellular distributions of these soluble scaffolds resemble Ssa2-GFP, which is found in the cytoplasm and puncta likely representing sites of EV biogenesis (Logan et al., 2024), whereas Tom70-GFP fluorescence was restricted to reticular structures reminiscent of mitochondria, as expected. Membrane spanning scaffolds fused with EGFP and mRuby2 are also found on intracellular puncta, but are absent from the cytoplasm, as expected, and are observed in foci on or adjacent to the plasma membrane. Their presence on or near the plasma membrane is consistent with their human cellular distributions (Seong et al., 2017), and patches observed are reminiscent of ectosomes, or lipid rafts, where some EVs are generated (Nydegger et al., 2006). In some cells, low levels of fluorescence are observed within the vacuole lumen, likely representing products of protein degradation.

**Figure 3.**
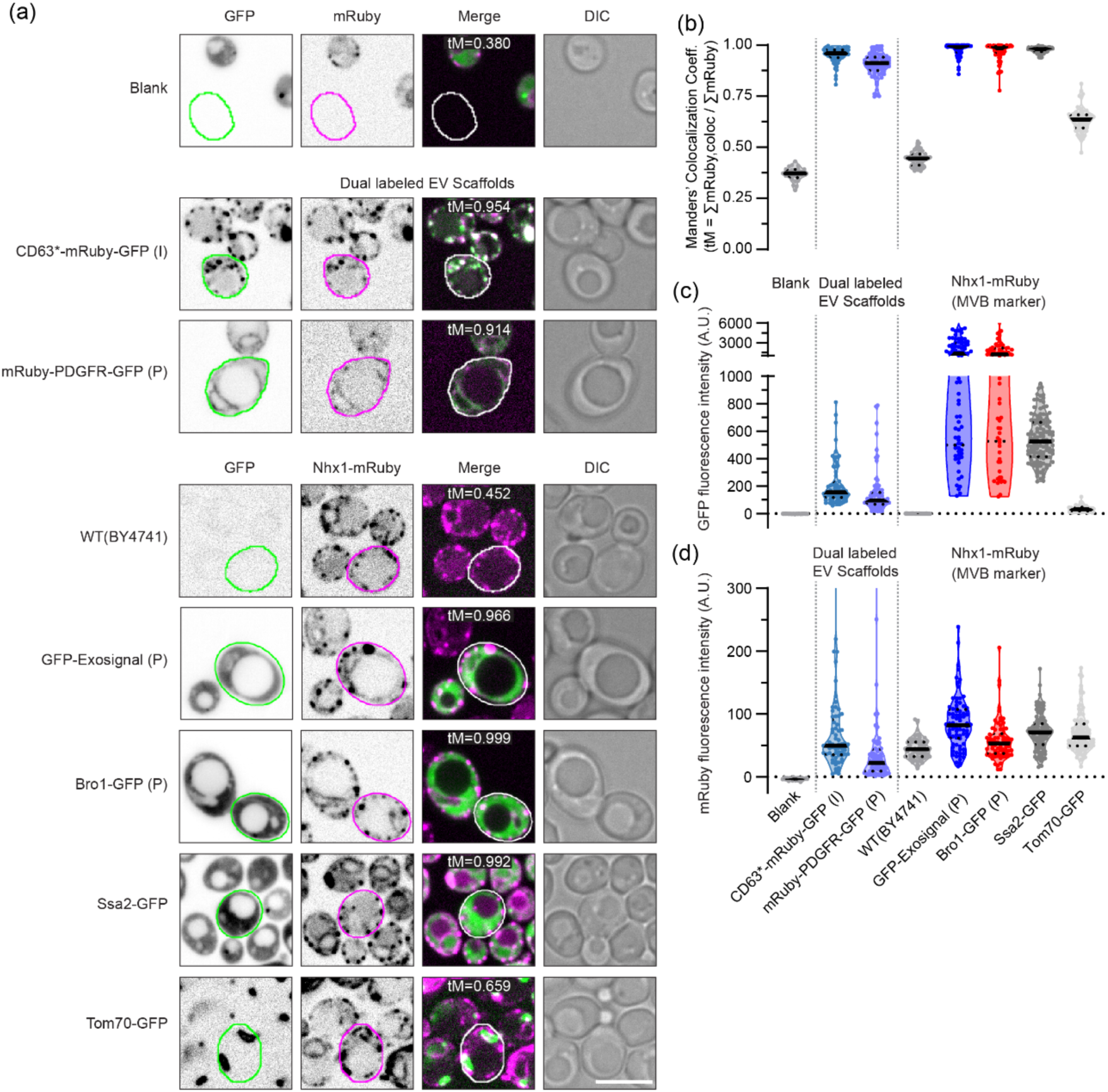
Candidate scaffold proteins colocalize with multivesicular bodies in *S. cerevisiae*. (a) Representative confocal microscopy images used for colocalization analysis. Yeast cells from all strains were imaged in GFP, mRuby (RFP), and DIC channels. A merge of the GFP and mRuby channels is also presented. Strains with a (P) indicate plasmid expression, strains with an (I) indicate genomic integration. Colocalization analysis was carried out on individual cells, using ROIs obtained via the YeastSpotter neural network or manually drawn on images. Colocalization analysis used the Fiji BIOP JACoP plugin, with the thresholded Manders’ colocalization coefficient (tM) as a readout. To determine the threshold and establish a lower bound for the tM, we analyzed the noise in micrographs for each strain by moving the cell ROIs to regions of the image without cells (Blank). Images presented show a representative ROI in green (GFP), magenta (mRuby), or white (Merge), with the tM for that ROI presented at the top of the Merge image. To establish an upper bound for the tM, dual labeled EV scaffolds with GFP and mRuby fused to the same protein were analyzed. To determine colocalization with MVBs, the MVB marker Nhx-1 fused to mRuby was genomically integrated into strains carrying EVclo proteins or controls labeled with just GFP. Fluorescent images presented are after background subtraction and brightness/contrast adjustment. Presented blank images are from the Ssa2-GFP strain, but blanks for each strain were quantified. Scale bar, 5 µm. (b) Thresholded Manders’ colocalization coefficients for strains analyzed for colocalization. The tM measures how much of the mRuby fluorescence intensity in that ROI (Ʃ mRuby) is found in pixels that are also above the threshold for GFP (Ʃ mRuby,coloc). Data presented are from 59-119 cell ROIs quantified from 1-30 images, plotted as violin plots with individual points shown, with solid lines marking the median and dotted lines marking the 1^st^ and 3^rd^ quartiles. Data presented are from a single biological replicate, with a second biological replicate showing identical trends. (c,d) Mean pixel intensities of the same ROIs presented in (b), for the GFP (c) and mRuby (d) channels. Mean pixel intensities were calculated after background subtraction, and the threshold used for colocalization analysis was subtracted from each value presented. Data are presented as violin plots with individual points shown, with solid lines marking the median and dotted lines marking the 1^st^ and 3^rd^ quartiles. Data presented are from a single biological replicate, with a second biological replicate showing identical trends. Data points presented are the average of 2 technical replicates, with points representing biological replicates and bars displaying S.E.M. P-values comparing 30°C vs. 42°C for each strain were calculated using Welch’s 2-tailed t-test.

Within eukaryotic cells, small EVs such as exosomes are biosynthesized at endosomal membranes to produce multivesicular bodies (MVBs) (van Niel et al., 2018). To assess whether scaffolds localize to MVBs, we fused mRuby2 to the C-terminus of Nhx1, a Na^+^(K^+^)/H^+^ exchanger exclusively embedded within MVB perimeter membranes (Karim & Brett, 2018), and integrated it into the genomes of strains containing single-labeled scaffolds fused to EGFP. We then visualized these live cells using confocal fluorescence microscopy (Figure 3a) and conducted colocalization analysis to determine if signal from Nhx1-mRuby2 and EGFP-labeled scaffolds overlap within cells (Figure 3b). As a lower bound for the colocalization metric (thresholded Manders’ Colocalization Coefficient, tM) we conducted analysis on blank regions of micrographs without cells, as a negative control we used wildtype cells expressing just Nhx1-mRuby2 without EGFP, and as positive controls for colocalization analysis we examined dual labeled scaffold proteins fused to both mRuby2 and EGFP (CD63*-mRuby-GFP, mRuby-PDGFR-GFP). Ssa2-GFP was previously shown to be packaged into exosomes (Logan et al., 2024) and we find that it colocalizes with the MVB marker Nhx1-mRuby2 as expected (Figure 3b), whereas the mitochondrial protein Tom70-GFP shows poor colocalization with Nhx1-mRuby2. Analysis of micrographs to quantify cellular EGFP fluorescence (Figure 3c) confirmed data obtained by flow cytometry (see Figure 2b), and cellular mRuby2 fluorescence is similar in all strains tested (Figure 3d), validating the colocalization analysis. Ultimately, we found that GFP-ExoSignal and Bro1-GFP both colocalized with Nhx1-mRuby2 at the same level as Ssa2-GFP, suggesting they are present at the sites of exosome biogenesis within cells.

### 3.4 | Human and yeast scaffolds are present in EVs released from *S. cerevisiae*

To determine if scaffold candidates are within extracellular vesicles, we first collected extracellular medium from yeast cultures resuspended in PBS and subjected to mild heat stress (42°C for 30 minutes) which triggers release of small EVs (Logan et al., 2024). We counted and sized the particles within EV fractions using nanoparticle tracking analysis (Figure 4a – c), and we visualized them using transmission electron microscopy (Figure 4d) to determine if over-expression of scaffold proteins alters biogenesis, release or morphology of EVs. We found that candidate scaffold expression did not significantly affect the diameter (Figure 4b), number (Figure 4c) or morphology (Figure 4d) of EV particles released from yeast cells. Notably, median particle diameter is near 142.5 nm for wild type and all strains tested, indicating that small EVs or exosomes are present in these preparations.

**Figure 4.**
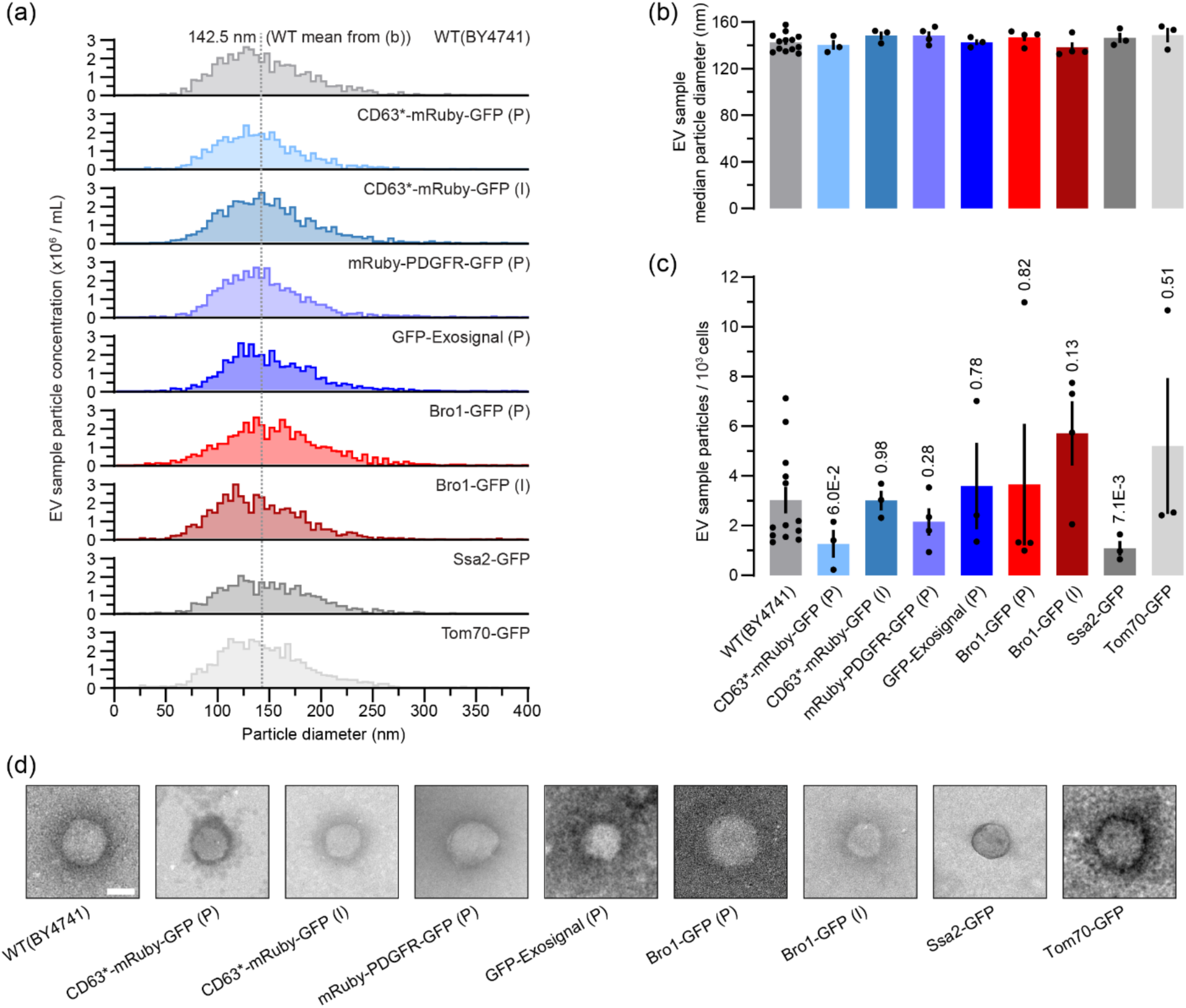
Scaffold protein expression does not alter EV production. (a) Representative nanoparticle tracking analysis (NTA) histograms. The vertical line at 142.5 nm, provided as a guide for the eye, is placed at the mean of the median particle diameters for WT(BY4741) presented in (b). (b) Median particle diameters from NTA. Each point represents a biological replicate, averaged over 2 technical replicates, bars display S.E.M. No strains are significantly different from WT(BY4741) by Welch’s 2-tailed t-test; WT(BY4741) vs. mRuby-PDGFR-GFP has the smallest p-value at 0.21. (c) Particles produced per thousand cells. Particle number was determined by NTA, cell number by OD and volume of culture. Each point represents a biological replicate, averaged over 2 technical replicates, bars display S.E.M. P-values presented are compared to WT(BY4741) by Welch’s 2-tailed t-test. (d) Representative negative-stained transmission electron micrographs. Scale bar, 50 nm.

To demonstrate that scaffold proteins are present in purified EV fractions, we measured EGFP fluorescence by fluorometry (Figure 5a). Of the six scaffold-containing strains tested, four produced EVs with GFP fluorescence significantly higher than background: Bro1-GFP (plasmid) > Bro1-GFP (integrated) > GFP-ExoSignal (plasmid) >> CD63*-mRuby-GFP (integrated), as did the positive control Ssa2-GFP. EVs collected from cells containing CD63*-mRuby-GFP (plasmid), mRuby-PDGFR-GFP (plasmid) or the negative control Tom70-GFP did not show EGFP signal over background.

**Figure 5.**
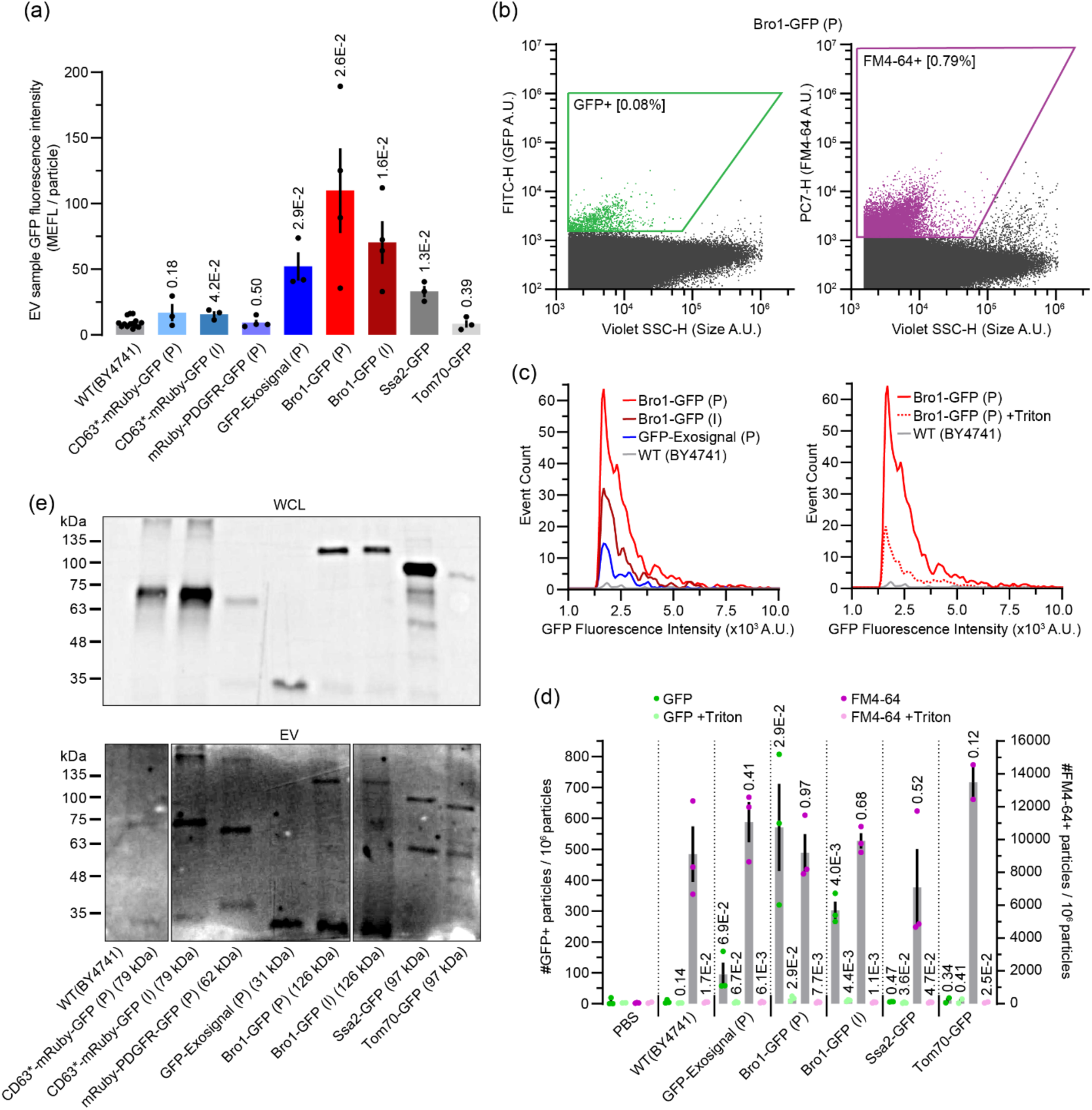
ExoSignal and Bro1 scaffolds drive GFP test cargo into *S. cerevisiae* EVs. (a) GFP fluorescence of EV samples, assayed using a plate reader. A standard curve of fluorescein was prepared in duplicate on the same plate to convert raw fluorescence to calibrated Molecules of Equivalent Fluorescein (MEFL) units. Particles numbers were obtained from nanoparticle tracking analysis (NTA). Dilution series of particles were assayed, obtaining multiple points with fluorescence above the limit of detection (LOD). Points presented represent biological replicates, with each point representing the average of all MEFL/particle values above the LOD, over two technical replicates. Bars display S.E.M. P-values presented are compared to WT(BY4741) via Welch’s 1-tailed t-test. (b) Representative nanoflow cytometry plots for one tested strain (Bro1-GFP (P)), with EVs stained using the membrane dye FM4-64. Data presented were gated for debris and singlets, GFP positive gate was determined using WT(BY4741) EV sample, FM4-64 positive gate was determined using PBS +FM4-64 sample. (c) (Left) Representative histograms of fluorescence intensity for GFP+ particles. Data are from nanoflow cytometry GFP+ gate. (Right) Representative histograms of fluorescence intensity for GFP+ particles before and after Triton detergent treatment using the same instrument, acquisition, and analysis parameters. (d) Compiled nanoflow cytometry data for promising EV scaffolds and controls before and after Triton treatment. The number of particles in GFP or FM4-64 positive gates were normalized to the total number of particles after gating for debris and singlets. Each point represents a biological replicate, bars display S.E.M. P-values for GFP (darker green) were compared to WT(BY4741) via Welch’s 1-tailed t-test, FM4-64 (darker purple) were compared to WT(BY4741) via Welch’s 2-tailed t-test, and +Triton conditions (lighter green and lighter purple) were compared to non-Triton treated for the same strain via Welch’s 1-tailed t-test. (e) Western blot analysis of whole cell lysate (top) or EV samples (bottom) for all strains tested, probed using antibodies raised against GFP. Predicted protein sizes are indicated after the strain name, and molecular weight marker locations are indicated along the left side of the blots.

To demonstrate that EGFP signal was present in lipid-bound particles (EVs), we conducted nano-flow cytometry on EV fractions stained with the membrane dye FM4-64 (Figure 5b – d). We limited our experiments to testing only Bro1-GFP (plasmid or integrated) and GFP-ExoSignal (plasmid) because robust EGFP signal was detected in EV samples (see Figure 5a), as well as positive (Ssa2-GFP) and negative (wild type no GFP, Tom70-GFP, PBS only) controls. Of all particles detected in these samples, approximately 1% were FM4-64-positive (e.g. Figure 5b and d) and this signal collapsed after adding the detergent Triton X-100 (Figure 5d), suggesting this fraction of particles is lipid-bound, representing EVs. Consistent with previous results, Bro1-GFP (plasmid) showed the highest proportion of GFP-positive particles (∼0.1% of total or 10% of the lipid-bound fraction) and highest particle fluorescent intensity detected (Figure 5b – d). Bro1-GFP (integrated) and GFP-ExoSignal fluorescence was also detected within particles, but Ssa2-GFP signal was presumably too low for detection over background (Figure 5d). Addition of Triton X-100 to all EV samples abolished EGFP signals, suggesting that tagged scaffolds are present within lipid-bound particles representing EVs (Figure 5c and d). To demonstrate that EGFP fusion proteins were intact within EV samples, we conducted Western blot analysis (Figure 5e). After confirming the presence of fusion proteins within whole cell lysates, we identify bands representing whole scaffold-GFP proteins in EV samples studied, similar to Ssa2-GFP (positive control). No bands were observed in whole cell lysates or EV samples from wild type cells (no GFP, negative control). Together, these results show that some yeast (Bro1-GFP) and human (GFP-ExoSignal) scaffolds are readily sorted into small EVs.

## 4 | DISCUSSION

Here we demonstrate a complete design–build–test–learn (DBTL) cycle for engineering yeast extracellular vesicles using principles of synthetic biology: To load desired cargoes into EVs, we (re)designed parts and modified a modular gene expression system to accommodate four CDS assembly strategies that fuse cargoes (EGFP, mRuby2) to potential EV scaffolds. Using this new EVclo system, we successfully built at least one construct for each strategy and tested CDSs encoded on plasmids or integrated into the genome. By examining fluorescent fusion protein expression in cells and EVs, we learned that – of the scaffold candidates tested – yeast Bro1 and human ExoSignal show potential for loading cargoes into yeast EVs. Supporting automated multi-cycle research workflows, this EVclo system can be used to screen additional scaffold candidates for improved EV cargo loading, and to replace fluorescent proteins with desired therapeutic cargoes. Moreover, this EVclo design allows easy CDS transfer to other cloning systems that support engineered EV biomanufacturing by other chassis organisms (e.g. bacteria, mesenchymal stem cells) (Li et al., 2025) (Yin et al., 2023) (Danilushkina et al., 2023).

This pilot study identifies the first scaffolds for EV cargo loading in *S. cerevisiae*. Of the candidates tested, Bro1showed highest accumulation in EVs. Loading seems to be selective as other proteins tested (Tom70, PDGFR. CD63) were not enriched in EVs (Figure 5). Bro1 is the yeast ortholog of human ALIX implicated in small EV (or exosome) biogenesis through interactions with syntenin-1, syndecan and tetraspanins like CD63 (Teng & Fussenegger, 2021). However, ALIX overexpression in human cells does not affect EV release (Sun et al., 2019).

This consistent with our results showing that overexpressing Bro1 alone in yeast does not affect EV titers (Figure 4c). It is worth noting that there are no clear orthologs of syntenin-1, syndecans or tetraspanins encoded in the *S. cerevisiae* genome, and human CD63 is not sorted into EVs when ectopically expressed in *S. cerevisiae* (Figure 5). Moreover, although ALIX and Bro1 share a conserved Bro1 domain presumed to mediate ESCRT-dependent cargo loading – a common function – the remainder of their peptide sequences have low homology (Tseng et al., 2022).

This suggests that an ALIX/syntenin/syndecan/tetrspanin-mediated EV biogenesis pathway may not be conserved in yeast and Bro1 may contribute to vesicle formation or be sorted into EVs using different mechanisms. Further resolving its role in this process is the focus of future studies.

ExoSignal is the only human scaffold tested that is enriched in EVs when ectopically expressed in *S. cerevisiae*. ExoSignal is a short peptide (VKKDQ-AEPLHR-KFERQ) containing KFERQ-like motifs from α -synuclein and ribonuclease A separated by a peptide spacer (Ferreira et al., 2022). These motifs are thought to interact with LAMP2A and/or HSP70 for sorting into small EVs released from cultured human ARPE-19 cells, and ExoSignal-tagged GFP has been found in circulating EVs in zebrafish larvae (Ferreira et al., 2022). *S. cerevisiae* does not possess a LAMP2A ortholog however our group has shown that the HSP70 ortholog Ssa2 is sorted into yeast EVs and use it as a positive control in this study (Logan et al., 2024). In *S. cerevisiae*, Hsp70 proteins bind to the Sup35 prion protein for disaggregation as well as propagation and transmission, which is in part mediated by EVs (Shen et al., 2024) (Liu et al., 2016) (Kabani & Melki, 2015). Given that ExoSignal contains a sequence for clearance and transmission of α-synuclein – a prion-like protein, we hypothesize that it may get sorted into EVs using an evolutionarily conserved HSP70-mediated pathway. Like Bro1, HSP70 is thought to interact with ESCRTs to mediate cargo loading into intralumenal vesicles or EVs (Tseng et al., 2022) (Sahu et al., 2011). Thus, we speculate that both scaffolds identified – Bro1 and ExoSignal –use a deeply conserved ESCRT-mediated mechanism to sort cargos into EVs. We will test these hypotheses in the future to better understand the mechanisms underlying small EV biogenesis using *S. cerevisiae* as a model.

Although the discovery of two EV scaffolds is promising, we recognize that, like other engineered EV platforms being developed (Li et al., 2023) (Rädler et al., 2023) (Herrmann et al., 2021), there is room for improvement: Our best scaffold, Bro1 (tagged to GFP and expressed from a plasmid), is detected within ∼550 particles in samples that contained 9,000 FM4-64-positive, detergent-sensitive particles when examined by nano-flow cytometry (Figure 5d). Thus, at best, we estimate that Bro1-GFP may be present in ∼6% of small EVs analyzed. This result permits us and others to initiate EV engineering projects that will include additional post-isolation bioprocessing steps to improve sample homogeneity (e.g. affinity purification (Bonner et al., 2024)). However, this EVclo system can be used to conduct wider screens (unbiased or focused, based on EV proteomics datasets; e.g. (Logan et al., 2024) (Mencher et al., 2020) (Zhao et al., 2019) focused on identifying more efficient lumenal scaffolds and the first membrane scaffolds in *S. cerevisiae*, as candidates tested in this pilot study, CD63 and PDGFR, were not detected in EVs (Figure 5). Thus, by establishing *S. cerevisiae* and EVclo as a platform to efficiently engineer EVs, we help move the field closer towards realizing the potential of EVs as a new drug delivery modality with many potential applications.

## AUTHOR CONTRIBUTIONS

J.B. – Conceptualization, Formal Analysis, Investigation, Methodology, Validation, Visualization, Writing – Original Draft Preparation; J.T. – Investigation, Validation; A.C.P. – Investigation, Validation; M.F. –Investigation, Validation; M. N.-S. – Investigation, Validation; C.L.B. – Conceptualization, Funding Acquisition, Methodology, Project Administration, Resources, Supervision, Visualization, Writing – Original Draft Preparation.

## ACKNOWLEDGEMENTS

We thank: Devina Singh for assistance with cloning; Hélène Pagé-Veillette, Nadim Tawil, and Laura Montermini at the Centre for Applied Nanomedicine at the McGill University Health Centre for assistance with nanoparticle tracking analysis and nano-flow cytometry; David Liu and Kelly Sears for assistance with transmission electron microscopy at the Facility for Electron Microscopy Research, McGill University (Montreal, Canada); and Christopher Law for help with fluorescence microscopy at the Centre for Microscopy and Cell Imaging, Concordia University. Flow cytometry was conducted in the Genome Foundry at Concordia University.

## DATA AVILABILITY STATEMENT

Data that support findings of this study are available from the corresponding author upon reasonable request.

## REFERENCES

1. Alberti, S., Gitler, A. D., & Lindquist, S. (2007). A suite of Gateway® cloning vectors for high-throughput genetic analysis in *Saccharomyces cerevisiae*. Yeast, 24(10), 913–919. 10.1002/yea.1502

2. Amin, S., Massoumi, H., Tewari, D., Roy, A., Chaudhuri, M., Jazayerli, C., Krishan, A., Singh, M., Soleimani, M., Karaca, E. E., Mirzaei, A., Guaiquil, V. H., Rosenblatt, M. I., Djalilian, A. R., & Jalilian, E. (2024). Cell Type-Specific Extracellular Vesicles and Their Impact on Health and Disease. International Journal of Molecular Sciences, 25(5). 10.3390/IJMS25052730

3. Baig, H., Fontanarossa, P., McLaughlin, J., Scott-Brown, J., Vaidyanathan, P., Gorochowski, T., Misirli, G., Beal, J., & Myers, C. (2021). Synthetic biology open language visual (SBOL visual) version 3.0. Journal of Integrative Bioinformatics, 18(3), 20210013. 10.1515/JIB-2021-0013

4. Bean, B. D. M., Whiteway, M., & Martin, V. J. J. (2022). The MyLO CRISPR-Cas9 toolkit: a markerless yeast localization and overexpression CRISPR-Cas9 toolkit. G3 Genes|Genomes|Genetics, 12(8). 10.1093/g3journal/jkac154

5. Bolte, S., & Cordelières, F. P. (2006). A guided tour into subcellular colocalization analysis in light microscopy. Journal of Microscopy, 224(3), 213–232. 10.1111/J.1365-2818.2006.01706.X

6. Bonner, S. E., van de Wakker, S. I., Phillips, W., Willms, E., Sluijter, J. P. G., Hill, A. F., Wood, M. J. A., & Vader, P. (2024). Scalable purification of extracellular vesicles with high yield and purity using multimodal flowthrough chromatography. Journal of Extracellular Biology, 3(2), e138. 10.1002/JEX2.138

7. Buchholz, K., & Collins, J. (2013). The roots - A short history of industrial microbiology and biotechnology. Applied Microbiology and Biotechnology, 97(9), 3747–3762. 10.1007/s00253-013-4768-2

8. Colombo, M., Raposo, G., & Théry, C. (2014). Biogenesis, secretion, and intercellular interactions of exosomes and other extracellular vesicles. Annual Review of Cell and Developmental Biology, 30, 255–289. 10.1146/annurev-cellbio-101512-122326

9. Danilushkina, A. A., Emene, C. C., Barlev, N. A., & Gomzikova, M. O. (2023). Strategies for Engineering of Extracellular Vesicles. International Journal of Molecular Sciences, 24(17), 13247. 10.3390/IJMS241713247

10. David, F., Davis, A. M., Gossing, M., Hayes, M. A., Romero, E., Scott, L. H., & Wigglesworth, M. J. (2021). A Perspective on Synthetic Biology in Drug Discovery and Development—Current Impact and Future Opportunities. SLAS Discovery, 26(5), 581–603. 10.1177/24725552211000669

11. Dooley, K., McConnell, R. E., Xu, K., Lewis, N. D., Haupt, S., Youniss, M. R., Martin, S., Sia, C. L., McCoy, C., Moniz, R. J., Burenkova, O., Sanchez-Salazar, J., Jang, S. C., Choi, B., Harrison, R. A., Houde, D., Burzyn, D., Leng, C., Kirwin, K., … Williams, D. E. (2021). A versatile platform for generating engineered extracellular vesicles with defined therapeutic properties. Molecular Therapy, 29(5), 1729–1743. 10.1016/J.YMTHE.2021.01.020

12. Endy, D. (2005). Foundations for engineering biology. Nature, 438(7067), 449–453. 10.1038/nature04342

13. Erana-Perez, Z., Igartua, M., Santos-Vizcaino, E., & Hernandez, R. M. (2024). Genetically engineered loaded extracellular vesicles for drug delivery. Trends in Pharmacological Sciences, 45(4), 350–365. 10.1016/j.tips.2024.02.006

14. Estes, S., Konstantinov, K., & Young, J. D. (2022). Manufactured extracellular vesicles as human therapeutics: challenges, advances, and opportunities. Current Opinion in Biotechnology, 77, 102776. 10.1016/J.COPBIO.2022.102776

15. Ferreira, J. V., Da Rosa Soares, A., Ramalho, J., Carvalho, C. M., Cardoso, M. H., Pintado, P., Carvalho, A. S., Beck, H. C., Matthiesen, R., Zuzarte, M., Girao, H., Van Niel, G., & Pereira, P. (2022). LAMP2A regulates the loading of proteins into exosomes. Science Advances, 8(12), 1140. 10.1126/SCIADV.ABM1140

16. Freemont, P. S. (2019). Synthetic biology industry: data-driven design is creating new opportunities in biotechnology. Emerging Topics in Life Sciences, 3(5), 651–657. 10.1042/ETLS20190040

17. Garcia, S., & Trinh, C. T. (2019). Modular design: Implementing proven engineering principles in biotechnology. Biotechnology Advances, 37(7), 107403. 10.1016/J.BIOTECHADV.2019.06.002

18. Garner, K. L. (2021). Principles of synthetic biology. Essays in Biochemistry, 65(5), 791. 10.1042/EBC20200059

19. Gill, S., Catchpole, R., & Forterre, P. (2019). Extracellular membrane vesicles in the three domains of life and beyond. FEMS Microbiology Reviews, 43(3), 273–303. 10.1093/FEMSRE/FUY042

20. Gryciuk, A., Milner-Krawczyk, M., Rogalska, M., Banach, A. K., Sitkiewicz, E., Bakun, M., Świadek, M. E., & Mierzejewska, J. (2025). Characteristics of Two *Saccharomyces cerevisiae* Strains and Their Extracellular Vesicles as New Candidates for Probiotics. Probiotics and Antimicrobial Proteins, 1–22. 10.1007/S12602-025-10624-0

21. Herrmann, I. K., Wood, M. J. A., & Fuhrmann, G. (2021). Extracellular vesicles as a next-generation drug delivery platform. Nature Nanotechnology, 16(7), 748–759. 10.1038/s41565-021-00931-2

22. Higuchi, A., Morishita, M., Nagata, R., Maruoka, K., Katsumi, H., & Yamamoto, A. (2023). Functional Characterization of Extracellular Vesicles from Baker’s Yeast *Saccharomyces Cerevisiae* as a Novel Vaccine Material for Immune Cell Maturation. Journal of Pharmaceutical Sciences, 112(2), 525–534. 10.1016/J.XPHS.2022.08.032

23. Huh, W. K., Falvo, J. V., Gerke, L. C., Carroll, A. S., Howson, R. W., Weissman, J. S., & O’Shea, E. K. (2003). Global analysis of protein localization in budding yeast. Nature, 425, 686–691. 10.1038/nature02026

24. Jafari, D., Shajari, S., Jafari, R., Mardi, N., Gomari, H., Ganji, F., Forouzandeh Moghadam, M., & Samadikuchaksaraei, A. (2020). Designer Exosomes: A New Platform for Biotechnology Therapeutics. Biodrugs, 34(5), 567. 10.1007/S40259-020-00434-X

25. Jeon, S., Sohn, Y. J., Lee, H., Park, J. Y., Kim, D., Lee, E. S., & Park, S. J. (2025). Recent advances in the Design-Build-Test-Learn (DBTL) cycle for systems metabolic engineering of Corynebacterium glutamicum. Journal of Microbiology, 63(3), e2501021. 10.71150/jm.2501021

26. Kabani, M., & Melki, R. (2015). Sup35p in its soluble and prion states is packaged inside extracellular vesicles. MBio, 6(4). 10.1128/MBIO.01017-15

27. Kalluri, R., & LeBleu, V. S. (2020). The biology, function, and biomedical applications of exosomes. Science, 367(6478). 10.1126/SCIENCE.AAU6977

28. Karim, M. A., & Brett, C. L. (2018). The Na^+^(K^+^)/H^+^ exchanger Nhx1 controls multivesicular body–vacuolar lysosome fusion. Molecular Biology of the Cell, 29(3), 317–325. 10.1091/MBC.E17-08-0496

29. Kelwick, R. J. R., Webb, A. J., & Freemont, P. S. (2024). Accelerating extracellular vesicle research and biotechnological applications using synthetic biology approaches. Extracellular Vesicle, 4, 100050. 10.1016/J.VESIC.2024.100050

30. Kojima, R., Bojar, D., Rizzi, G., Hamri, G. C. El, El-Baba, M. D., Saxena, P., Ausländer, S., Tan, K. R., & Fussenegger, M. (2018). Designer exosomes produced by implanted cells intracerebrally deliver therapeutic cargo for Parkinson’s disease treatment. Nature Communications, 9(1). 10.1038/s41467-018-03733-8

31. Kumar, M. A., Baba, S. K., Sadida, H. Q., Marzooqi, S. Al, Jerobin, J., Altemani, F. H., Algehainy, N., Alanazi, M. A., Abou-Samra, A. B., Kumar, R., Al-Shabeeb Akil, A. S., Macha, M. A., Mir, R., & Bhat, A. A. (2024). Extracellular vesicles as tools and targets in therapy for diseases. Signal Transduction and Targeted Therapy, 9. 10.1038/s41392-024-01735-1

32. Lawson, C. E., Harcombe, W. R., Hatzenpichler, R., Lindemann, S. R., Löffler, F. E., O’Malley, M. A., García Martín, H., Pfleger, B. F., Raskin, L., Venturelli, O. S., Weissbrodt, D. G., Noguera, D. R., & McMahon, K. D. (2019). Common principles and best practices for engineering microbiomes. Nature Reviews Microbiology, 17(12), 725–741. 10.1038/s41579-019-0255-9

33. Lee, M. E., DeLoache, W. C., Cervantes, B., & Dueber, J. E. (2015). A Highly Characterized Yeast Toolkit for Modular, Multipart Assembly. ACS Synthetic Biology, 4(9), 975–986. 10.1021/SB500366V

34. Li, G., Chen, T., Dahlman, J., Eniola-Adefeso, L., Ghiran, I. C., Kurre, P., Lam, W. A., Lang, J. K., Marbán, E., Martín, P., Momma, S., Moos, M., Nelson, D. J., Raffai, R. L., Ren, X., Sluijter, J. P. G., Stott, S. L., Vunjak-Novakovic, G., Walker, N. D., … Sundd, P. (2023). Current challenges and future directions for engineering extracellular vesicles for heart, lung, blood and sleep diseases. Journal of Extracellular Vesicles, 12(2), 12305. 10.1002/jev2.12305

35. Li, Q., Chen, X., Xie, J., & Nie, S. (2025). Engineered Bacterial Extracellular Vesicles: Developments, Challenges, and Opportunities. Engineering, 54(16), 291–307. 10.1016/j.eng.2025.06.042

36. Liebana-Jordan, M., Brotons, B., Falcon-Perez, J. M., & Gonzalez, E. (2021). Extracellular Vesicles in the Fungi Kingdom. International Journal of Molecular Sciences, 22(13). 10.3390/IJMS22137221

37. Liu, S., Hossinger, A., Hofmann, J. P., Denner, P., & Vorberg, I. M. (2016). Horizontal transmission of cytosolic sup35 prions by extracellular vesicles. MBio, 7(4), 915–931. 10.1128/MBIO.00915-16

38. Logan, C. J., Staton, C. C., Oliver, J. T., Bouffard, J., David, T., Kazmirchuk, D., Magi, M., & Brett, C. L. (2024). Thermotolerance in *S. cerevisiae* as a model to study extracellular vesicle biology. Journal of Extracellular Vesicles, 13(5). 10.1002/jev2.12431

39. Lu, A. X., Zarin, T., Hsu, I. S., & Moses, A. M. (2019). YeastSpotter: accurate and parameter-free web segmentation for microscopy images of yeast cells. Bioinformatics, 35(21). 10.1093/bioinformatics/btz402

40. Ludwig, A. K., & Giebel, B. (2012). Exosomes: Small vesicles participating in intercellular communication. The International Journal of Biochemistry & Cell Biology, 44(1), 11–15. 10.1016/J.BIOCEL.2011.10.005

41. Ma, Y., Dong, S., Grippin, A. J., Teng, L., Lee, A. S., Kim, B. Y. S., & Jiang, W. (2025). Engineering therapeutical extracellular vesicles for clinical translation. Trends in Biotechnology, 43(1), 61–82. 10.1016/J.TIBTECH.2024.08.007

42. Malcl, K., Watts, E., Roberts, T. M., Auxillos, J. Y., Nowrouzi, B., Boll, H. O., Nascimento, C. Z. S. Do, Andreou, A., Vegh, P., Donovan, S., Fragkoudis, R., Panke, S., Wallace, E., Elfick, A., & Rios-Solis, L. (2022). Standardization of Synthetic Biology Tools and Assembly Methods for *Saccharomyces cerevisiae* and Emerging Yeast Species. ACS Synthetic Biology, 11(8), 2527–2547. 10.1021/ACSSYNBIO.1C00442

43. Mathieu, M., Martin-Jaular, L., Lavieu, G., & Théry, C. (2019). Specificities of secretion and uptake of exosomes and other extracellular vesicles for cell-to-cell communication. Nature Cell Biology, 21(1), 9–17. 10.1038/s41556-018-0250-9

44. Mencher, A., Morales, P., Valero, E., Tronchoni, J., Patil, K. R., & Gonzalez, R. (2020). Proteomic characterization of extracellular vesicles produced by several wine yeast species. Microbial Biotechnology, 13(5), 1581–1596. 10.1111/1751-7915.13614

45. Mentkowski, K. I., Snitzer, J. D., Rusnak, S., & Lang, J. K. (2018). Therapeutic Potential of Engineered Extracellular Vesicles. The AAPS Journal, 20(3). 10.1208/S12248-018-0211-Z

46. Mizenko, R. R., Feaver, M., Bozkurt, B. T., Lowe, N., Nguyen, B., Huang, K. W., Wang, A., & Carney, R. P. (2024). A critical systematic review of extracellular vesicle clinical trials. Journal of Extracellular Vesicles, 13(10), e12510. 10.1002/JEV2.12510

47. Nydegger, S., Khurana, S., Krementsov, D. N., Foti, M., & Thali, M. (2006). Mapping of tetraspanin-enriched microdomains that can function as gateways for HIV-1. Journal of Cell Biology, 173(5), 795–807. 10.1083/JCB.200508165

48. Oliveira, D. L., Nakayasu, E. S., Joffe, L. S., Guimarães, A. J., Sobreira, T. J. P., Nosanchuk, J. D., Cordero, R. J. B., Frases, S., Casadevall, A., Almeida, I. C., Nimrichter, L., & Rodrigues, M. L. (2010). Characterization of Yeast Extracellular Vesicles: Evidence for the Participation of Different Pathways of Cellular Traffic in Vesicle Biogenesis. PLOS ONE, 5(6), e11113. 10.1371/JOURNAL.PONE.0011113

49. Paganini, C., Capasso Palmiero, U., Pocsfalvi, G., Touzet, N., Bongiovanni, A., & Arosio, P. (2019). Scalable Production and Isolation of Extracellular Vesicles: Available Sources and Lessons from Current Industrial Bioprocesses. Biotechnology Journal, 14(10), 1800528. 10.1002/BIOT.201800528

50. Rädler, J., Gupta, D., Zickler, A., & Andaloussi, S. EL. (2023). Exploiting the biogenesis of extracellular vesicles for bioengineering and therapeutic cargo loading. Molecular Therapy, 31(5), 1231–1250. 10.1016/j.ymthe.2023.02.013

51. Rodrigues, M., Fan, J., Lyon, C., Wan, M., & Hu, Y. (2018). Role of Extracellular Vesicles in Viral and Bacterial Infections: Pathogenesis, Diagnostics, and Therapeutics. Theranostics, 8(10), 2709. 10.7150/THNO.20576

52. Sahu, R., Kaushik, S., Clement, C. C., Cannizzo, E. S., Scharf, B., Follenzi, A., Potolicchio, I., Nieves, E., Cuervo, A. M., & Santambrogio, L. (2011). Microautophagy of Cytosolic Proteins by Late Endosomes. Developmental Cell, 20(1), 131–139. 10.1016/j.devcel.2010.12.003

53. Seong, J., Huang, M., Sim, K. M., Kim, H., & Wang, Y. (2017). FRET-based Visualization of PDGF Receptor Activation at Membrane Microdomains. Scientific Reports, 7(1). 10.1038/s41598-017-01789-y

54. Shen, C. H. H., Komi, Y., Nakagawa, Y., Kamatari, Y. O., Nomura, T., Kimura, H., Shida, T., Burke, J., Tamai, S., Ishida, Y., & Tanaka, M. (2024). Exposed Hsp70-binding site impacts yeast Sup35 prion disaggregation and propagation. Proceedings of the National Academy of Sciences of the United States of America, 121(51), e2318162121. 10.1073/pnas.2318162121

55. Stickney, Z., Losacco, J., McDevitt, S., Zhang, Z., & Lu, B. (2016). Development of exosome surface display technology in living human cells. Biochemical and Biophysical Research Communications, 472(1), 53–59. 10.1016/J.BBRC.2016.02.058

56. Stranford, D. M., Simons, L. M., Berman, K. E., Cheng, L., DiBiase, B. N., Hung, M. E., Lucks, J. B., Hultquist, J. F., Leonard, J. N., & Biomed Eng, N. (2024). Bioengineering multifunctional extracellular vesicles for targeted delivery of biologics to T cells. Nat Biomed Eng, 8(4), 397–414. 10.1038/s41551-023-01142-x

57. Sun, R., Liu, Y., Lu, M., Ding, Q., Wang, P., Zhang, H., Tian, X., Lu, P., Meng, D., Sun, N., Xiang, M., & Chen, S. (2019). ALIX increases protein content and protective function of iPSC-derived exosomes. Journal of Molecular Medicine, 97(6), 829–844. 10.1007/S00109-019-01767-Z

58. Teng, F., & Fussenegger, M. (2021). Shedding Light on Extracellular Vesicle Biogenesis and Bioengineering. Advanced Science, 8(1), 2003505. 10.1002/ADVS.202003505

59. Tkach, M., & Théry, C. (2016). Communication by Extracellular Vesicles: Where We Are and Where We Need to Go. Cell, 164(6), 1226–1232. 10.1016/j.cell.2016.01.043

60. Tseng, C. C., Piper, R. C., & Katzmann, D. J. (2022). Bro1 family proteins harmonize cargo sorting with vesicle formation. BioEssays, 44(8), 2100276. 10.1002/BIES.202100276

61. Van Delen, M., Derdelinckx, J., Wouters, K., Nelissen, I., & Cools, N. (2024). A systematic review and meta-analysis of clinical trials assessing safety and efficacy of human extracellular vesicle-based therapy. Journal of Extracellular Vesicles, 13(7), e12458. 10.1002/JEV2.12458

62. van Niel, G., Carter, D. R. F., Clayton, A., Lambert, D. W., Raposo, G., & Vader, P. (2022). Challenges and directions in studying cell–cell communication by extracellular vesicles. Nature Reviews Molecular Cell Biology, 23(5), 369–382. 10.1038/s41580-022-00460-3

63. van Niel, G., D’Angelo, G., & Raposo, G. (2018). Shedding light on the cell biology of extracellular vesicles. Nature Reviews Molecular Cell Biology, 19(4), 213–228. 10.1038/nrm.2017.125

64. Vignoni, A., Boada, Y., Boada Acosta, L., Andreu-Vilarroig, C., Alarcón, I., Requena, A, & Picó, J. (2019). Fluorescence calibration and color equivalence for quantitative synthetic biology. IFAC-PapersOnLine, 52(26), 129–134. 10.1016/j.ifacol.2019.12.247

65. Welsh, J. A., Goberdhan, D. C. I., O’Driscoll, L., Buzas, E. I., Blenkiron, C., Bussolati, B., Cai, H., Di Vizio, D., Driedonks, T. A. P., Erdbrügger, U., Falcon-Perez, J. M., Fu, Q. L., Hill, A. F., Lenassi, M., Lim, S. K., Mahoney, M. ỹ. G., Mohanty, S., Möller, A., Nieuwland, R., … Zubair, H. (2024). Minimal information for studies of extracellular vesicles (MISEV2023): From basic to advanced approaches. Journal of Extracellular Vesicles, 13(2), e12404. 10.1002/JEV2.12404

66. Whitford, C. M., Cruz-Morales, P., Keasling, J. D., & Weber, T. (2021). The Design-Build-Test-Learn cycle for metabolic engineering of *Streptomycetes*. Essays in Biochemistry, 65(2), 261–275. 10.1042/EBC20200132

67. Woith, E., Fuhrmann, G., & Melzig, M. F. (2019). Extracellular Vesicles—Connecting Kingdoms. International Journal of Molecular Sciences, 20(22), 5695. 10.3390/IJMS20225695

68. Xu, G., Jin, J., Fu, Z., Wang, G., Lei, X., Xu, J., & Wang, J. (2025). Extracellular vesicle-based drug overview: research landscape, quality control and nonclinical evaluation strategies. Signal Transduction and Targeted Therapy, 10(1). 10.1038/s41392-025-02312-w

69. Yáñez-Mó, M., Siljander, P. R. M., Andreu, Z., Zavec, A. B., Borràs, F. E., Buzas, E. I., Buzas, K., Casal, E., Cappello, F., Carvalho, J., Colás, E., Cordeiro-Da Silva, A., Fais, S., Falcon-Perez, J. M., Ghobrial, I. M., Giebel, B., Gimona, M., Graner, M., Gursel, I., … De Wever, O. (2015). Biological properties of extracellular vesicles and their physiological functions. Journal of Extracellular Vesicles, 4(1), 1–60. 10.3402/JEV.V4.27066

70. Yin, T., Liu, Y., Ji, W., Zhuang, J., Chen, X., Gong, B., Chu, J., Liang, W., Gao, J., & Yin, Y. (2023). Engineered mesenchymal stem cell-derived extracellular vesicles: A state-of-the-art multifunctional weapon against Alzheimer’s disease. Theranostics, 13(4), 1264–1285. 10.7150/thno.81860

71. Yuan, M., Ma, W., Liu, B., Zou, X., Huang, B., Tian, X., Jin, Y., Zheng, N., Wu, Z., & Wang, Y. (2024). Delivery of therapeutic RNA by extracellular vesicles derived from *Saccharomyces cerevisiae* for medicine applications. Journal of Pharmaceutical Sciences, 113(12), 3574–3585. 10.1016/J.XPHS.2024.10.035

72. Zhao, K., Bleackley, M., Chisanga, D., Gangoda, L., Fonseka, P., Liem, M., Kalra, H., Al Saffar, H., Keerthikumar, S., Ang, C. S., Adda, C. G., Jiang, L., Yap, K., Poon, I. K., Lock, P., Bulone, V., Anderson, M., & Mathivanan, S. (2019). Extracellular vesicles secreted by *Saccharomyces cerevisiae* are involved in cell wall remodeling. Communications Biology, 2(1). 10.1038/s42003-019-0538-8

