## Supporting Materials for "Engineering *S. cerevisiae* extracellular vesicles using synthetic biology"

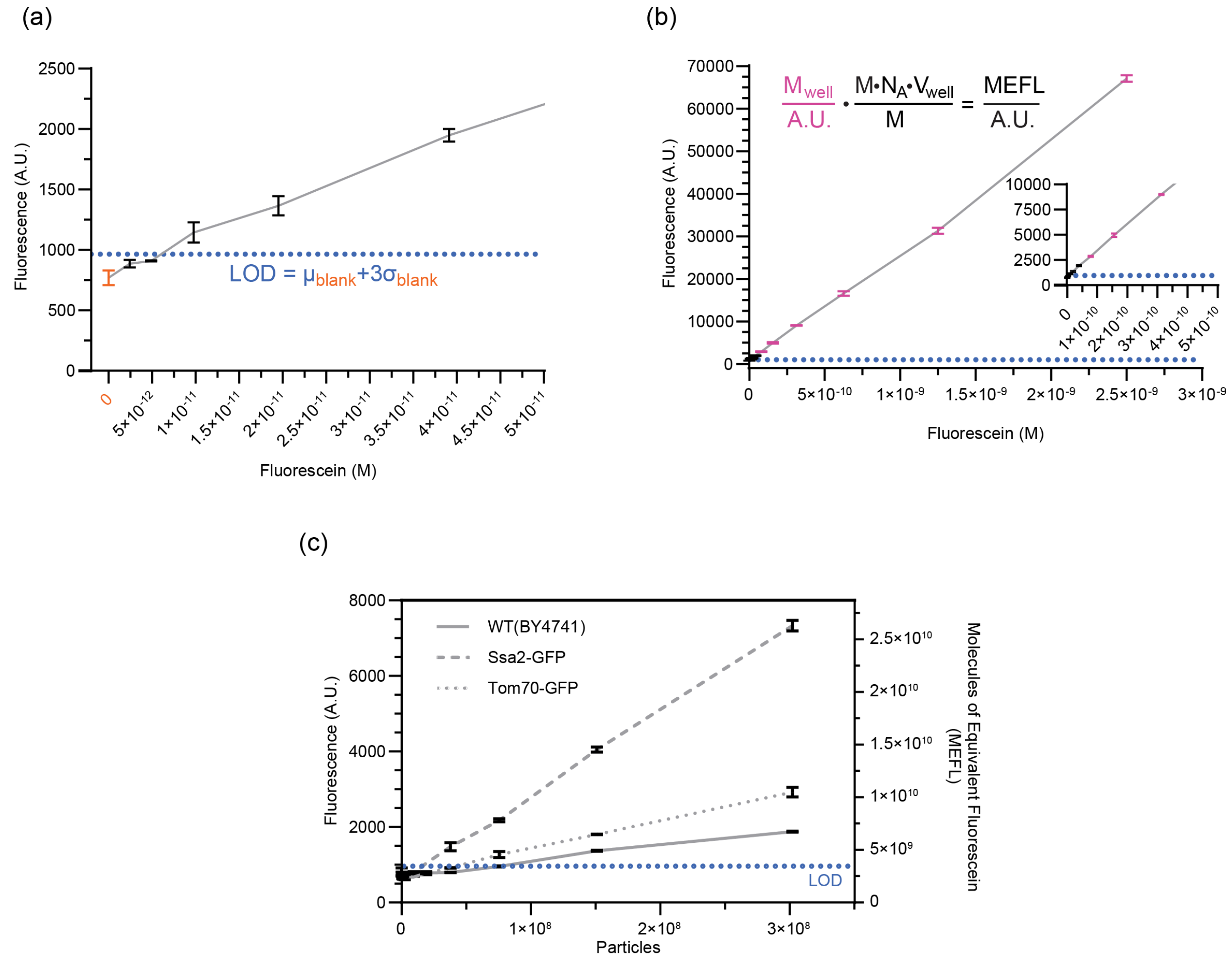

**Supplemental Figure 1 (previous page)**

Molecules of Equivalent Fluorescein (MEFL) calibration.

For plate reader assays, calibration curves using a fluorescein standard were prepared in duplicate on the same plate as samples. Bars on the curves display the S.E.M. over the duplicates for each point. Fluorescein dilutions were prepared in the same ultrapure PBS as samples, starting at 2.5 nM and doing 10 1:2 serial dilutions, covering 3 orders of magnitude for fluorescein concentration. The last well of each row contained just PBS as blanks. (a) shows the lowest fluorescein concentrations, reaching the noise floor of the plate reader, and (b) shows the highest concentrations with the inset showing intermediate concentrations. To calculate the MEFL/A.U. conversion factor, first the limit of detection (LOD) was determined by calculating the mean and standard deviation of the fluorescence from the blank wells (a, orange), with LOD (blue dotted lines) defined as the mean + 3 standard deviations. To remove background from the fluorescent signals, this LOD value was subtracted from all fluorescence values, and points below the LOD were excluded from further analysis. Given differences in the fluorescent spectra between our calibrant fluorescein and sample GFP, we applied a spectral correction factor to the background subtracted fluorescence. These background subtracted and spectrally corrected fluorescence values provided the denominator for the $M_{well}$/A.U. term (magenta) in the equation in (b), with the known concentration of the well providing the numerator. The second term in this equation provides a conversion factor to molecules of fluorescein, with the numerator calculating the molecules per well by multiplying the initial molarity (M), Avogadro's number ($N_A$), and the well volume ($V_{well}$), and the denominator providing the same initial molarity as the numerator to cancel units with the molarity of the well in the first term. Multiplying the two terms gives molecules of fluorescein per arbitrary unit of fluorescence. We found empirically that the 6 highest fluorescein concentrations (magenta) provided the most stable MEFL/A.U. conversion factor, so just these points were used and averaged to generate a single MEFL/A.U. conversion factor. (c) To calibrate the fluorescence from samples, raw fluorescence values below the LOD were discarded, the remaining values were background subtracted and spectrally corrected, and the fluorescence values in arbitrary units were multiplied by the MEFL/A.U. conversion factor.

(a)

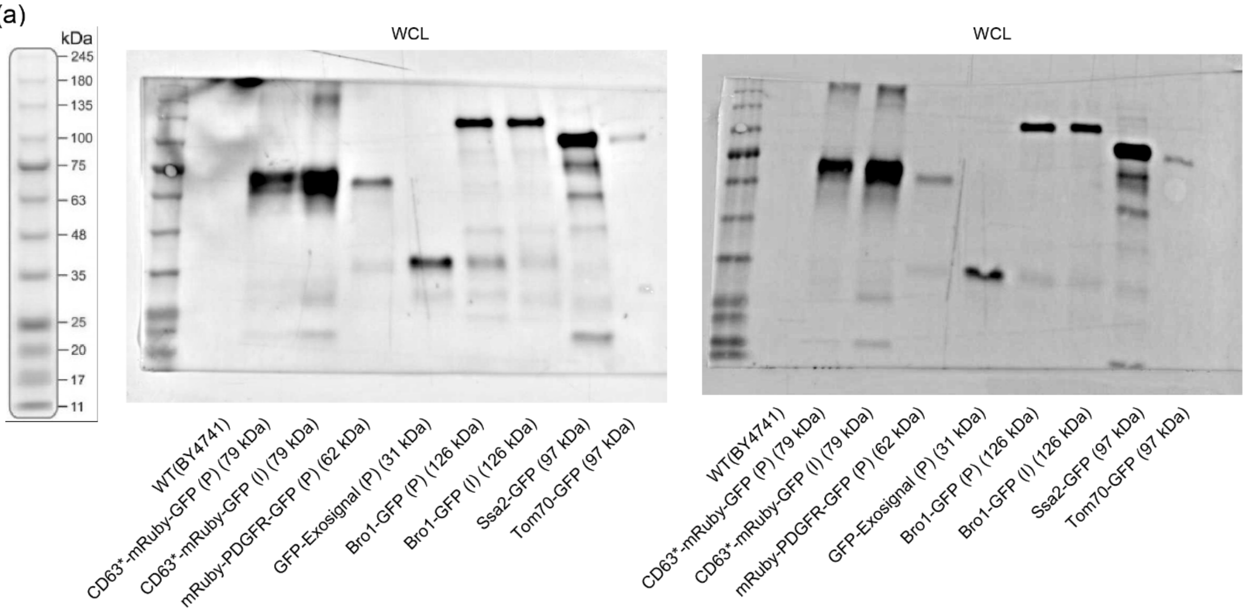


(b)

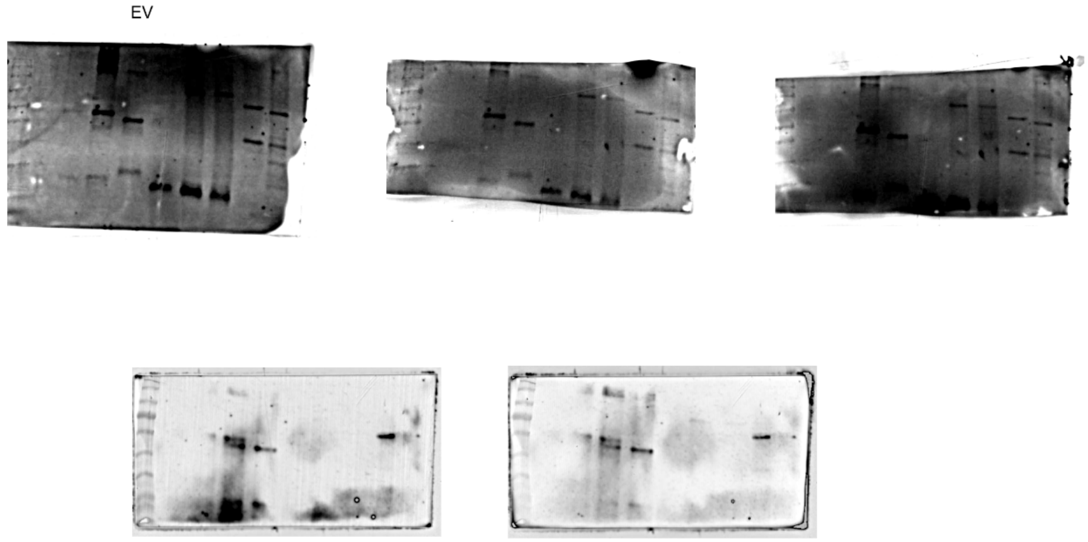


**Supplemental Figure 2**

Uncropped Western blot images.

(a) 2 different blots of whole cell lysate. (b) 2 different blots of EV samples, imaged at multiple exposure times. Ladder and lane order are the same as (a).
